# Direct reprogramming of human somatic cells into neuron-like cells involves a transition through a transient intermediate state

**DOI:** 10.64898/2026.05.14.725118

**Authors:** Carlos Bueno, Marta Martinez-Morga, Francisco J. Rodriguez-Lozano, David García-Bernal, Salvador Martínez, José M. Moraleda, Miguel Blanquer

**Affiliations:** Department of Haematology, Hospital Clínico Universitario Virgen de la Arrixaca, Instituto Murciano de Investigación Biosanitaria Pascual Parrilla (IMIB), University of Murcia, Murcia, 30003, Spain; Instituto de Neurociencias de Alicante (UMH-CSIC), Universidad Miguel Hernandez, San Juan, Alicante, 03550, Spain; Department of Dermatology, Stomatology, Radiology and Physical Medicine, Morales Meseguer Hospital, Faculty of Medicine, IMIB-Arrixaca, University of Murcia, Murcia, Spain; Biochemistry, Molecular Biology and Immunology Department, University of Murcia, Faculty of Medicine, Murcia, 30100, Spain; Center of Biomedical Network Research on Mental Health (CIBERSAM), Instituto de Salud Carlos III, 28029, Madrid, Spain; Alicante Institute for Health and Biomedical Research (ISABIAL), Alicante, 03010, Spain

**Keywords:** direct conversion, reprograming, transdifferentiation, neuronal differentiation, neurogenesis, mesenchymal, stem cells, somatic cells

## Abstract

**Background:** Direct conversion of human somatic cells into functional neurons could offer a faster way to generate patient-specific neurons for use in regenerative medicine, disease modelling, and drug development. Although it has been reported that neuronal direct reprogramming bypasses the intermediate pluripotent state, no reports have included time-lapse experiments, potentially overlooking transient intermediate states. Recent studies have shown that the conversion of human mesenchymal stromal cells (hMSCs) into neuron-like cells involves a transition through a transient intermediate state. Therefore, further research is needed to fully understand the process by which human somatic cells can become neurons without cell division. In this study we investigates whether direct neuronal reprogramming of human bone marrow-derived MSC (hBM-MSCs), dental pulp-derived MSC (hDP-MSCs), and adult human dermal fibroblasts (HDFa), involves a transient intermediate state, and sought to further validate the neuronal identity of hMSC-derived induced neurons.

**Methods:** In this study, we conducted time-lapse experiments to observe the transformation of hBM-MSCs, hDP-MSCs and HDFa, into neurons using a small-molecule-based direct reprogramming protocol. Cellular and ultrastructural changes were further characterized by confocal and electron microscopy.

**Results:** Direct conversion of hBM-MSCs, hDP-MSCs and HDFa into neuron-like cells occurred rapidly and in absence of cell division. Time-lapse analyses revealed that reprogramming proceeds through a transient intermediate state characterized by distinct morphological changes and dynamic nuclear remodelling. Furthermore, we found that neuron-like cells derived from hBM-MSCs and hDP-MSCs exhibit neuronal polarization, expressed specific neuronal and synaptic markers, formed interconnected cellular networks, and exhibited functional plasticity, providing further evidence that hMSCs can become functional neurons.

**Conclusions:** This study provides clear evidence that the direct neuronal reprogramming process involves a transition through an intermediate, transient state. Our findings also provide further evidence that hMSCs can become functional neurons. In summary, our work provides new insights into the direct neuronal reprogramming process, which is essential for advancing both developmental biology and regenerative medicine.

## Introduction

Access to healthy or diseased human neural tissue for research purposes remains a major limitation not only to advancing our understanding of the cellular and molecular mechanisms underlying neurogenesis and neurodegeneration, but also to creating autologous cells to replace damaged neurons in neurological disorders. For this reason, an enormous effort has been made over the last few decades to derive neurons from somatic cells isolated from various human tissue sources [1–4].

The conversion of somatic cells to functional neurons has been achieved using different starting cell populations and applying various techniques, such as indirect reprogramming [5, 6, 7], direct differentiation [2, 8] and direct reprogramming (often termed transdifferentiation) [4, 9–19]. These approaches rely on the use, either alone or in combination, of overexpression of transcription factors, non-coding RNAs, and small molecules. These in vitro-generated neuronal cells express specific neuronal markers, form synapses, and generate action potentials [10, 12, 15, 18]. Furthermore, they have been observed to functionally mature and integrate into the recipient’s neural network following transplantation [20, 21].

The emergence of technologies that enable the conversion of adult human somatic cells into neuronal cells not only offers a unique opportunity to investigate key aspects of central nervous system function, development, and disease at the cellular level, but also holds significant promise for their application in cell replacement therapies [16]. Despite rapid and significant advances in cell conversion technologies, the underlying mechanisms remain poorly understood, and this process is still considered inefficient, poorly controlled, and complex [16, 17, 19, 22, 23].

Therefore, more research is needed to understand the mechanisms governing cellular conversion and improve its efficiency, with the aim of harnessing their potential in regenerative medicine, disease modelling, and drug discovery.

Beyond their clinical implications, studies on lineage conversion and reprogramming have clearly demonstrated that somatic cell identity is not permanently fixed, but rather remains plastic and interconvertible, representing one of the most conceptual advances in modern developmental biology [24, 25]. Accordingly, future research into lineage conversion and reprogramming is essential for better understanding the mechanisms governing cell fate decisions in the field of developmental biology.

Previous studies have reported that direct reprogramming of somatic cells into neurons bypasses pluripotency and canonical developmental intermediates [9–19, 23, 24]. Mouse embryonic fibroblasts (MEFs) and human adult fibroblasts (HDFa) are the most commonly used starting cell types for direct reprogramming to generate functional neurons. However, it is important to note that these reports do not include time-lapse experiments [9–19], which allow for direct and continuous visualization of cellular fate conversion, thereby overcoming the limitations of fixed-cell analysis, which often overlooks short-lived transient states.

In our laboratory, we focus on the direct reprogramming (or transdifferentiation) of human mesenchymal stromal cells (hMSCs) into neuronal cells using small molecule-based protocols [26–30]. In previous publications, we showed that the conversion of hMSCs into neuron-like cells involves a transition through a transient intermediate state [27–30]. During this cellular state, the cells adopt a spherical morphology, and the nuclei are highly dynamic: they move through the cytoplasm, change shape and position, and even form cellular protrusions as they attempt to make contact with surrounding cells, likely to sense their microenvironment [29, 30]. We also observed that intermediate cells can preserve their spherical shape for several days without assuming new fates, or they can adopt alternative fates; they gradually adopted a neuron-like morphology through active neurite extension or re-differentiated back to the mesenchymal state. Furthermore, we found that intermediate cells undergo rapid and repeated lineages transitions without cell division [29].

In this study, we conducted long-term live cell imaging to observe the conversion of hMSCs derived from bone marrow (hBM-MSCs) and dental pulp (hDP-MSCs), as well as HDFa, into neurons. Time-lapse analysis allowed us not only to determine whether the direct reprogramming of somatic cells into neurons involves passage through a transient intermediate state, but also to provide additional evidence supporting the neuronal identity of hMSC-derived neurons.

## Methods

### Isolation and culture of hBM-MSCs, hDP-MSCs and HDFa

A standard protocol for the isolation and expansion of hBM-MSCs and hDP-MSCs was used as previously described [28–31]. Bone marrow aspirates were obtained by percutaneous direct aspiration from the iliac crest of 5 healthy volunteers at University Hospital Virgen de la Arrixaca (Murcia, Spain). Bone marrow was collected with sodium heparin (20 U/ml; Sigma-Aldrich) followed by Ficoll density gradient-based separation by centrifugation at 540 g for 20 min. After, the mononuclear cell fractions from different subjects were collected and mixed, washed twice with Ca2+/Mg2+-free phosphate buffered saline (PBS; Lonza). The cell suspension was plated into six-well multiwell plates (BD Falcon) and incubated at 37 °C in 5% CO2. In a previous publication, we showed that hBM-MSCs did not express hematopoietic lineage markers such as CD45, CD14, CD34 and CD20 and were positive for CD90, CD105, and CD73, thereby demonstrating a characteristic immunophenotype of hMSCs [29].

hDP-MSCs were harvested from the dental pulp of extracted impacted third molars from donors aged 16 to 22 years (n = 10). Human dental pulp (hDP) was digested with type I collagenase (3 mg/ml; Sigma-Aldrich) and dispase II (4 mg/L; Sigma-Aldrich) in Dulbecco’s Modified Eagle Medium (DMEM; Gibco) for 1 h at 37 °C. The reaction was stopped by the addition of DMEM. hDP samples derived from different subjects were pooled together to obtain single-cell suspension by passing the cells through a 70 μm strainer (BD Falcon). The cell suspension was plated into six-well multiwell plates (BD Falcon) and incubated at 37 °C in 5% CO2. In a previous publication, we showed that hDP -MSCs did not express hematopoietic lineage markers such as CD45, CD14, CD34 and CD20 and were positive for CD90, CD105, and CD73, thereby demonstrating a characteristic immunophenotype of hMSCs [31].

Adult human dermal fibroblasts (P10858 HDFa from Innoprot, Derio, Spain), hBM-MSCs and hDP-MSCs were cultured under standard conditions in serum-containing media (designated as basal media), composed of Dulbecco’s Modified Eagle Medium (DMEM; Gibco) supplemented with 10% fetal bovine serum (FBS; Lonza), 2 mM L-glutamine (Thermo Fisher Scientific), nonessential amino acid solution (Sigma-Aldrich) and 1% penicillin/streptomycin (Thermo Fisher Scientific). hBM-MSCs, hDP-MSCs and HDFa were cultured at 37°C and 5% CO2. Culture media were renewed every 3 days. All studies were performed using hMSCs expanded within culture passages 3-4.

### Time-lapse microscopy of hBM-MSCs, hDP-MSCs and HDFa cultured in neural induction media

We used 48-well plates for live-cell imaging. A vertical and a horizontal line were scored with a utility knife on the bottom (rear) of the plastic plate well. These physical marks serve as focal reference points that appear as sharp, dark shadows in phase-contrast images. Photographs were taken at the intersections of the lines (as shown in Fig.16).

To induce neural differentiation of hBM-MSCs, hDP-MSCs, cells were allowed to adhere overnight to plates coated with collagen IV (Sigma-Aldrich). Basal media was removed the following day, and the cells were cultured for 2 days in serum-free media (designated as the neural basal media) consisting of a 1:1 mixture of Dulbecco’s modified Eagle’s medium/F12 (DMEM/F12 Glutamax; Gibco) and Neurobasal Plus medium (Gibco) supplemented with 1% N2-supplement (Gibco) and 2% B-27 Plus-supplement (Gibco). On day 3, the cells were cultured in neural induction media consisting of neural basal media supplemented with Forskolin (20 μM; MedChemExpress), dibutyryl cAMP (200 μM; STEMCELL Technologies), L-Ascorbic acid (200 μM; MedChemExpress), retinoic acid (10 μM; MedChemExpress), BDNF (10 ng/mL; PeproTech), GDNF (10 ng/mL; PeproTech), NT-3 (10 ng/mL; PeproTech), CNTF (10 ng/mL; PeproTech) and bNGF (10 ng/mL; PeproTech). hBM-MSCs and hDP-MSCs were cultured with or without glass circle coverslips placed on top of them.

To induce neural differentiation of HDFa, cells were allowed to adhere overnight to plates coated with collagen IV (Sigma-Aldrich). Basal media was removed the following day, and the cells were cultured for 2 days in serum-free media (designated as the neural basal media) consisting of a 1:1 mixture of Dulbecco’s modified Eagle’s medium/F12 (DMEM/F12 Glutamax, Gibco) and Neurobasal Plus medium (Gibco) supplemented with 0.5% N2-supplement (Gibco) and 1% B-27 Plus-supplement (Gibco). On day 3, the cells were cultured in neural induction media consisting of neural basal media supplemented with dibutyryl-cAMP (1 mM; STEMCELL Technologies), L-Ascorbic acid (200 μM; MedChemExpress), BDNF (10 ng/mL; PeproTech), GDNF (10 ng /mL; PeproTech), NT-3 (10 ng/mL; PeproTech), CNTF (10 ng/mL; PeproTech) and bNGF (10 ng/mL; PeproTech). HDFa were cultured without glass circle coverslips placed on top of them.

Time-lapse analysis was performed using an inverted Leica DM IRB microscope. We conducted time-lapse microscopy during the first 12 days following the direct addition of the neural induction medium to the cells. Serial live-cell images were captured using a 10X objective every 24 hours. Data are representative of ten independent experiments.

### Immunocytochemistry

A standard immunocytochemical protocol was used as previously described [26, 30]. hBM-MSCs, hDP-MSCs and HDFa were seeded on glass circle coverslips coated with collagen IV (Sigma-Aldrich). The coverslips were then inverted, with the cell side facing down, and maintained in a neural induction medium. Cells were rinsed with PBS and fixed in freshly prepared 4% paraformaldehyde (PFA; Sigma-Aldrich). Fixed cells were blocked for 2 h in PBS containing 10% normal horse serum (Gibco) and 0.25% Triton X-100 (Sigma-Aldrich) and incubated overnight at 4°C with antibodies against β-III-tubulin (1:500; BioLegend), Doublecortin (DCX, 1/200; Abcam), GABA (1/200; Sigma-Aldrich), GAD65/67 (1/200; Sigma-Aldrich), GAP-43 (1/200; Abcam), GAT-1 (1/200; Chemicon), MAP-2 (1/200; Chemicon), Pvalb (1/200; Abcam), PSD-95 (1/200; Abcam), SHANK3 (1/500; Novusbio), Synapsin I (1/200; Novusbio), Synaptophysin (1/200; Abcam) and TAU-1 (1/200; Chemicon) in PBS containing 1% normal horse serum and 0.25% Triton X-100. The next day, the cells were rinsed and incubated with secondary antibodies conjugated with Alexa Fluor® 488 (anti-mouse or anti-rabbit; 1:500, Molecular Probes) and Alexa Fluor® 594 (anti-mouse or anti-rabbit; 1:500, Molecular Probes). Cell nuclei were counterstained with DAPI (0.2 mg/ml in PBS; Molecular Probes). Alexa Fluor 488® phalloidin (Molecular Probes) was used to selectively stain F-actin. Data are representative of ten independent experiments per condition.

### Scanning electron microscopy

HDFa were seeded on glass circle coverslips coated with collagen IV. The coverslips were then inverted, with the cell side facing down, and maintained in a neural induction medium. Cells were treated with fixative for 20 minutes. Coverslips were postfixed in 1% osmium tetroxide for 1 hour and dehydrated in graded ethanol washes. The coverslips were allowed to dry at a conventional critical point and were then coated with gold-palladium sputter-coated. Coverslips were view on a field emission scanning electron microscope (FESEM ApreoS).

### Images and Data Analyses

Photographs of visible and fluorescent stained samples were obtained using an inverted Leica DM IRB microscope equipped with a digital camera Leica DFC350FX (Nussloch) or a Leica Stellaris confocal laser scanning microscope. We used Leica Application Suite X software for confocal image analysis. Photoshop software was used to improve the visibility of fluorescence images without altering the underlying data.

## Results

### Chemical conversion of human somatic cells to neuron-like cells

As mentioned above, the aim of this study is to observe the conversion of hBM-MSCs and hDP-MSCs, as well as HDFa, into neurons, to determine whether the direct reprogramming of human somatic cells into neurons actually involves passage through a transient intermediate state. In addition, we examine whether neuron-like cells derived from human somatic cells behave similarly to primary neurons in culture, establishing neuronal polarity, expressing specific neuronal and synaptic markers, forming interconnected networks and developing electrical excitability.

Time-lapse imaging is the best method for validating neuronal conversion, as it allows for direct and continuous visualization of the conversion of the cellular fate, bypassing the limitations of fixed cell analysis, which often overlooks short-lived transient states. In this study, we were unable to use an automated time-lapse microscope to image live human somatic cells. Therefore, we have had to adapt multi-well plates to image the same regions at different time points (as described in Materials and Methods), which made it impossible to create time-lapse videos of cell conversion.

Despite the technical limitations involved in conducting a time-lapse experiment using conventional microscopy, the observations revealed that when hBM-MSCs were exposed to neural induction medium, they reshaped from a flat to a spherical morphology. We then observed the emergence of new neurite-like extensions sprouting directly from the cell bodies of these rounded cells. Subsequently, these cells gradually acquired a neuron-like morphology and established connections with one another through various types of interactions (Fig. 1A). Although it is possible to observe the early stages of neuronal differentiation of hBM-MSCs, time-lapse imaging also revealed that a large percentage of neuron-like cells derived from hBM-MSCs die during the first week of differentiation (Fig.1B), which makes it impossible to analyse long-term neuronal differentiation.

**Fig. 1.**
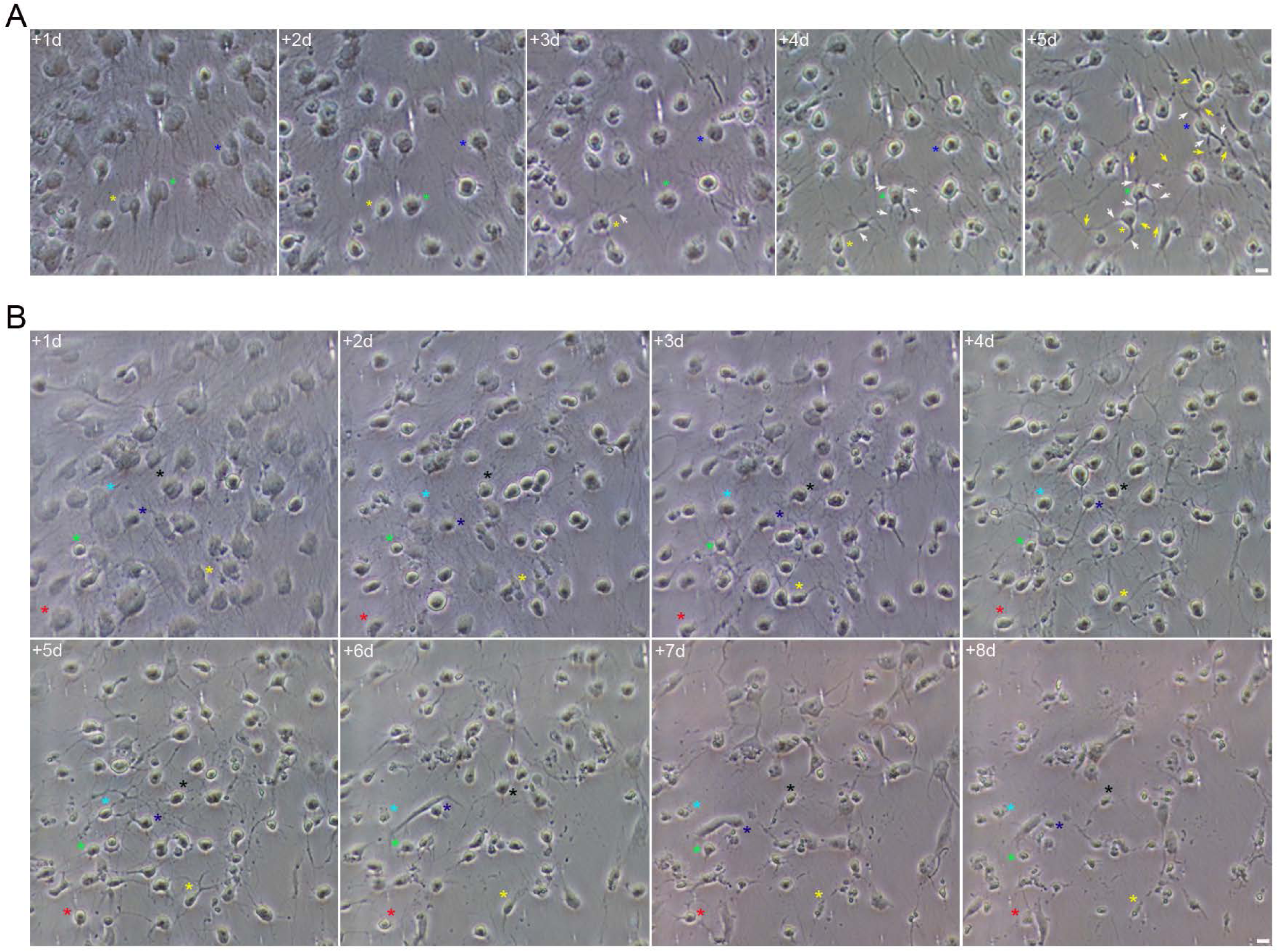
Chemical conversion of hBM-MSCs to neuron-like cells. (**A**) Time-lapse imaging revealed that when hBM-MSCs were exposed to neural induction medium, they reshaped from a flat to a spherical morphology. We then observed the emergence of new neurite-like extensions sprouting directly from the cell bodies of these rounded cells (white arrows). Subsequently, these cells gradually acquired a neuron-like morphology and established connections with one another through various types of interactions (yellow arrows). (**B**) However, we observed that a large percentage of neuron-like cells derived from hBM-MSCs die during the first week of differentiation, which makes it impossible to analyse long-term neuronal differentiation. Scale bar: 10 μm. The number at the top indicates the time since the time-lapse image began. Elapsed time is displayed in days.

It is important to note that it is well-documented that a large percentage of neurons in culture, particularly those derived from stem cells or primary tissue, die during the first week to two weeks of differentiation [1, 32, 33]. Neuron cell cultures are highly susceptible to early death due to high metabolic activity, rapid accumulation of neurotoxic substances and the extreme sensitivity to environmental changes including the light exposure used in live-cell imaging [32–34]. To determine whether the imaging process was the cause of the observed cell death, we used non-imaging controls. We observed that both groups are dying at a similar rate. Therefore, the observed cell death is not merely a consequence of the imaging process itself.

To minimise cell death and improve viability during neural differentiation, we have tested some different small-molecule-based direct reprogramming protocols that have been used to convert somatic cells into induced neurons [19]. Surprisingly, we have discovered that the most effective way to improve the survival of neuron-like cells derived from hBM-MSCs during neuronal differentiation is to place a glass coverslip over the cells. This method enables neuron-like cells derived from hBM-MSCs to be cultured for at least 24 days, with improved viability. We observed that neuron-like cells derived from hBM-MSCs cultured under a coverslip can survive while showing morphological maturation and the formation of interconnected networks (Fig. 2A). However, time-lapse imaging studies indicate that it is only possible to obtain images of living neuron-like cells derived from hBM-MSCs, cultured under a coverslip for up to 12 days, in which progressive morphological maturation and the formation of interconnected networks can be observed (Fig. 3). Therefore, neuron-like cells derived from hBM-MSCs subjected to daily imaging have shown lower viability compared to non-imaged controls. As noted above, we did not use an automated time-lapse microscope capable of maintaining a strict and stable environment. Therefore, the decreased viability of neuron-like cells derived from hBM-MSCs cultured under a coverslip during neural differentiation may result from prolonged exposure of the cells to environmental and light factors outside their normal physiological range.

**Fig. 2.**
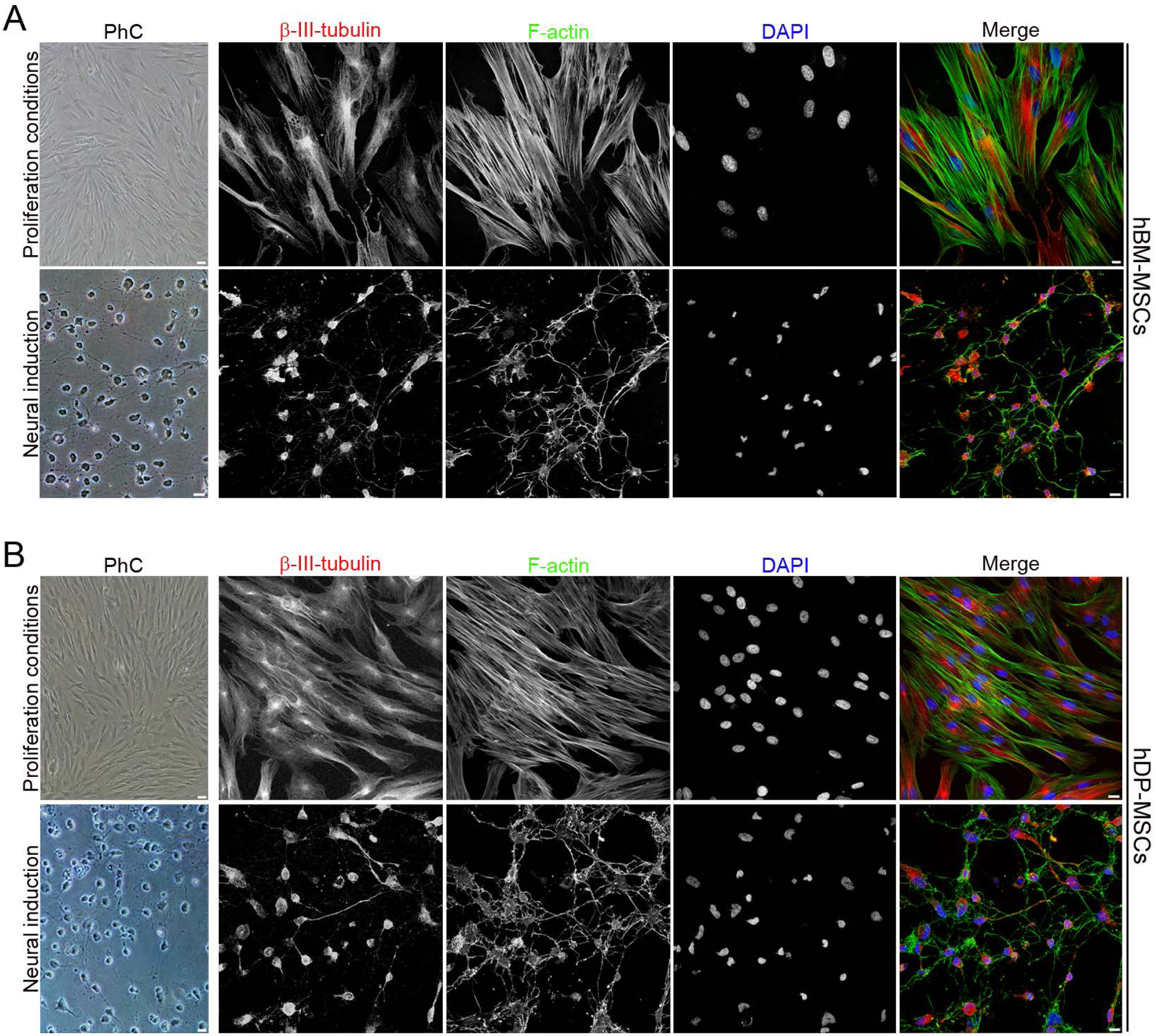
Placing a coverslip over hMSCs improves their viability during neural differentiation. Phase-contrast microscopy and cytoskeletal analysis show that placing a coverslip directly one cultured hBM-MSCs (**A**) and hDP-MSCs (**B**) improves cell survival, allowing for morphological maturation and the formation of interconnected networks. Scale bar: 10 μm. PhC: Phase-contrast photomicrographs.

**Fig. 3.**
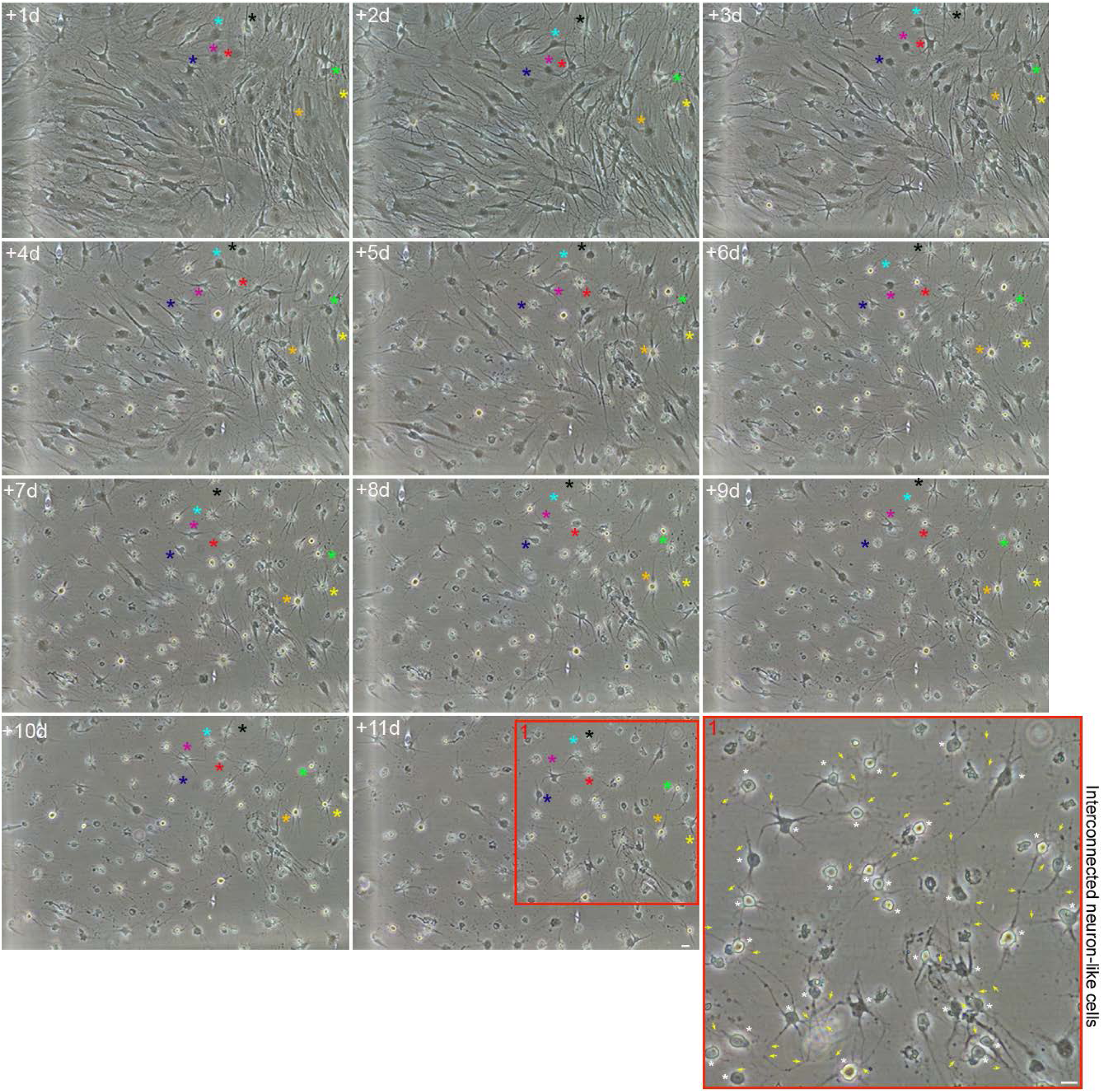
Neuron-like cells derived from hBM-MSCs cultured under a coverslip form interconnected networks. Time-lapse imaging studies indicate that it is possible to obtain images of living neuron-like cells derived from hBM-MSCs cultured under a coverslip for up to 12 days, in which progressive morphological maturation and the formation of interconnected networks can be observed. The white asterisks indicate interconnected neuron-like cells, and the yellow arrows show the connection points between these cells. Scale bar: 10 μm. The number at the top indicates the time since the time-lapse image began. Elapsed time is displayed in days.

To determine whether the coverslip method improves survival of neuron-like cells derived from other human somatic cells, we tested hDP-MSCs under the same neural induction conditions. Time-lapse imaging revealed that when hDP-MSCs not cultured under a coverslip were exposed to neural induction medium, they also reshaped from a flat to a spherical morphology. We then observed the emergence of new neurite-like extensions sprouting directly from the cell bodies of these rounded cells. Subsequently, these cells gradually acquired a neuron-like morphology and established connections with one another through various types of interactions (Fig. 4A).

**Fig. 4.**
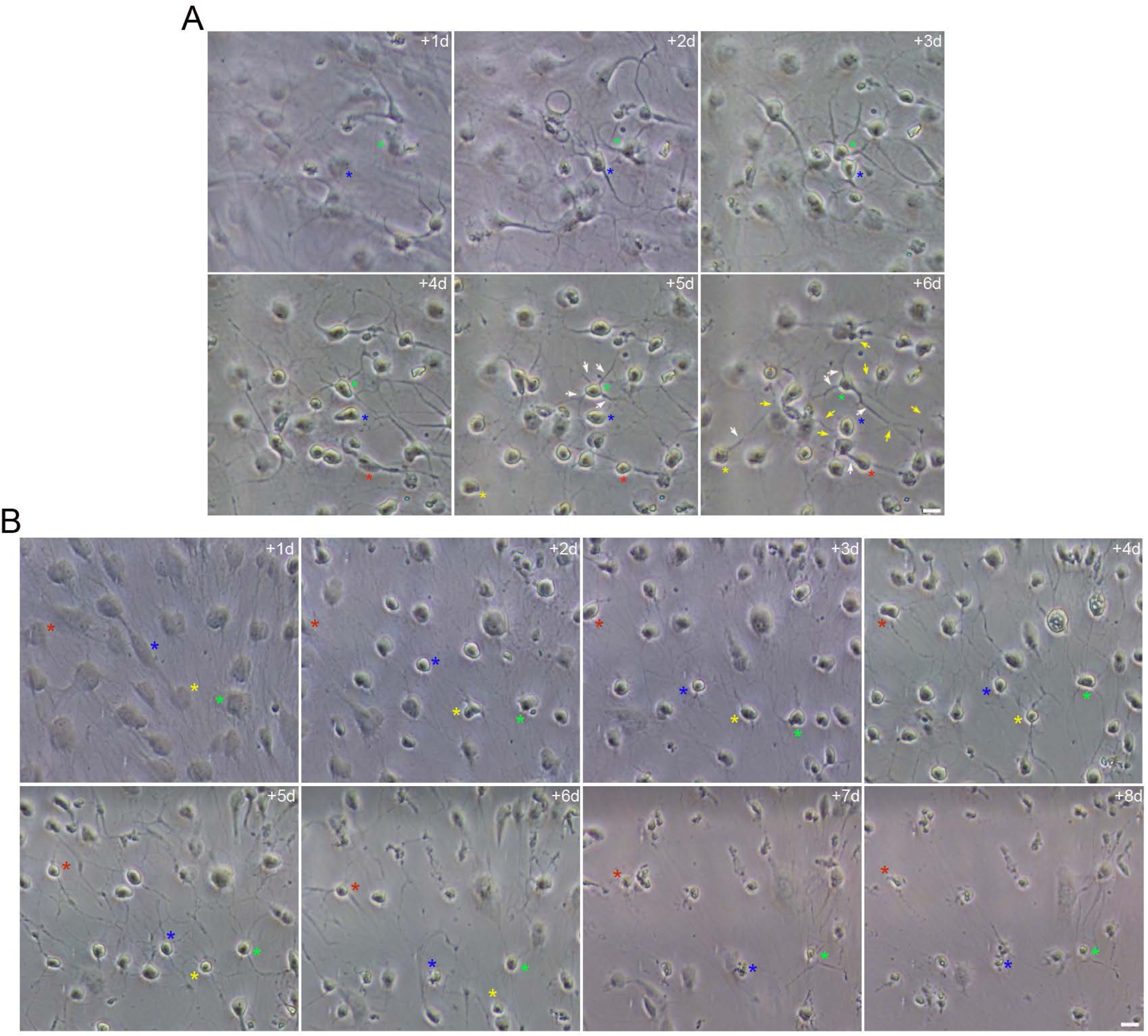
Chemical conversion of hDP-MSCs to neuron-like cells not cultured under a coverslip. (**A**) Time-lapse imaging revealed that when hDP-MSCs were exposed to neural induction medium, they reshaped from a flat to a spherical morphology. We then observed the emergence of new neurite-like extensions sprouting directly from the cell bodies of these rounded cells (white arrows). Subsequently, these cells gradually acquired a neuronal-like morphology and established connections with one another through various types of interactions (yellow arrows). (**B**) As with hBM-MSCs, we observed that a large percentage of neuron-like cells derived from hDP-MSCs die during the first week of differentiation. Scale bar: 10 μm. The number at the top indicates the time since the time-lapse image began. Elapsed time is displayed in days.

As with hBM-MSCs, we observed that a large percentage of neuron-like cells derived from hDP-MSCs die during the first week of differentiation (Fig. 4B), and that the viability rate of these cells, which were monitored daily, was similar to that of the control groups from which no images were obtained. However, when neuron-like cells derived from hDP-MSCs that have not been monitored daily using imaging techniques are cultured under a coverslip, they can survive for at least 24 days, while exhibiting morphological maturation and the formation of interconnected networks (Fig. 2B). As with hBM-MSCs, we also observed that neuron-like cells derived from hDP-MSCs subjected to daily imaging have shown lower viability compared to non-imaged controls. Time-lapse imaging studies indicate that it is only possible to obtain images of living neuron-like cells derived from hDP-MSCs, cultured under a coverslip for up to 12 days, in which progressive morphological maturation and the formation of interconnected networks can be observed (Fig. 5).

**Fig. 5.**
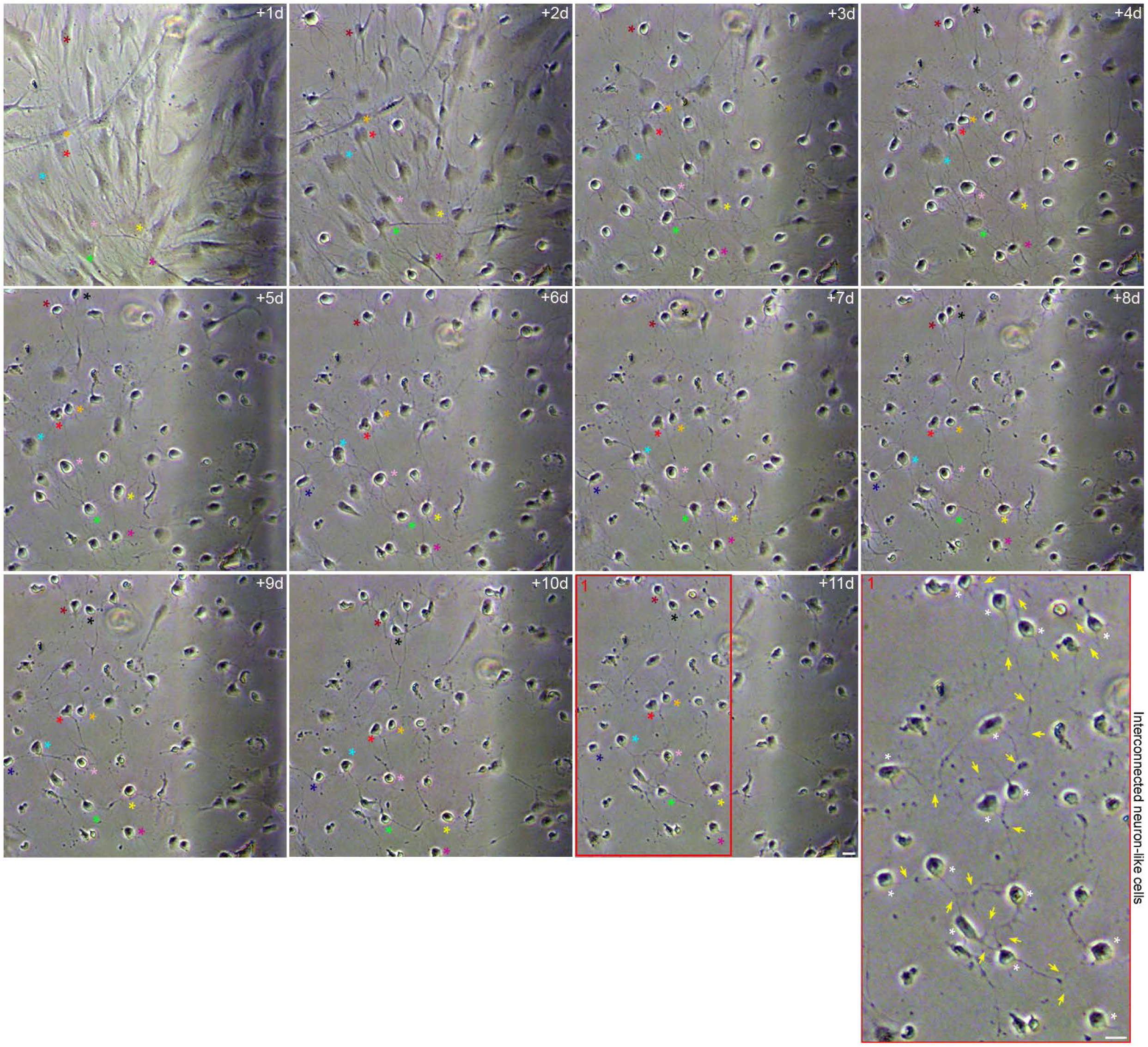
Neuron-like cells derived from hDP-MSCs cultured under a coverslip form interconnected networks. Time-lapse imaging studies indicate that it is possible to obtain images of living neuron-like cells derived from hDP-MSCs cultured under a coverslip for up to 12 days, in which progressive morphological maturation and the formation of interconnected networks can be observed. The white asterisks indicate interconnected neuron-like cells, and the yellow arrows show the connection points between these cells. Scale bar: 10 μm. The number at the top indicates the time since the time-lapse image began. Elapsed time is displayed in days.

It is important to note that no cell division is observed during the conversion of hBM-MSCs and hDP-MSCs into neuron-like cells (Figs. 1,3, 4, 5, 6, 12, 13, 14, 15). These findings are consistent with previous studies that reported that the direct reprogramming of human somatic cells into neuronal cells also occurs without cell division [9, 12, 15].

Our results provide evidence that placing a glass coverslip directly over the cells improves the long-term survival of neuron-like cells derived from human somatic cells. Further research is needed to understand the mechanisms underlying this phenomenon. This work also highlights that conducting time-lapse experiments using conventional microscopy without environmental controls severely compromises cell survival and hinders the analysis of long-term neuronal differentiation. Future research should be conducted using In-incubator microscopes equipped with time-lapse technology, to improve the analysis of long-term neuronal differentiation.

### Evidence of the neuronal identity of neuron**-**like cells derived from hMSCs

The authenticity of human-derived neurons, whether generated from iPS cells or through direct reprogramming, is validated by comparing their morphological, molecular, and functional characteristics with those described in primary human neural cultures and in vivo neurons. This process involves demonstrating that human-derived neurons exhibit neuronal polarization, express specific neuronal and synaptic markers, form interconnected networks, and display electrical excitability. The morphological progression of cultured rodent neurons follows a well-characterized developmental sequence in vitro, progressing from a simple, spherical cell body to a complex, interconnected network [35, 36]. The process of neuronal polarization begins with the formation of multiple minor neurites followed by the rapid elongation of one neurite to become the axon, while the others develop into dendrites. Subsequently, neurites branch extensively and form synapses, transforming the individual cells into a dense, interconnected network [35, 36].

Despite the technical limitations of conducting a time-lapse experiment using conventional microscopy without environmental controls, the observations obtained indicate that hBM-MSCs and hDP-MSCs display a morphological development sequence highly similar to those reported before in dissociated cell cultures derived from rodent brain (Figs. 6, 12, 13, 14). Time-lapse imaging reveals the emergence of new neurite-like extensions sprouting directly from the cell bodies of these newly hBM-MSCs and hDP-MSCs derived-round cells. Subsequently, we observed the formation of cells that exhibit a long neurite-like extension and multiple short neurite-like extensions extending from the opposite side of the cell body, resulting in cells that resemble neuronal morphology (Fig. 6). Immunocytochemical analysis confirm the neuronal morphology and structural changes observed in time-lapse microscopy (Fig. 7A). Cytoskeletal protein class III β-tubulin and F-actin staining shown that neuron-like cells develop distinct dendrite and axon-like processes. Class III β-tubulin forms a nearly continuous core along the length of the dendrite shaft and F-actin is largely concentrated in the dendritic periphery and protrusion, similar to the compartmentalized organization of the neuronal cytoskeleton described in dendrites [37] (Fig. 7B). Neuron-like cells derived from hBM-MSCs and hDP-MSCs also display long neurite-like extension with various types of axonal branch-like structures that mimic neurons [38], including terminal arborization (Fig. 7C), bifurcation (Fig. 7D), and collateral formation (Fig. 7E). We also observed that neuron-like cells produce long neurite-like extension featuring growth cones at their tips, with a spatial organisation similar to that described in neurons [39]. The central domain of the growth cone contains class III β-tubulin microtubules, and the peripheral domain comprises radial F-actin bundles (Fig. 7F). Immunocytochemical analysis also revealed that neuron-like cells derived from hBM-MSCs (Fig. 8) and hDP-MSCs (Fig. 9) express neural markers such as MAP-2, Tau-1, βIII-Tubulin, SHANK3, DCX, GAP-43, Pvalb, GABA, GAD65/67 and GAT-1.

**Fig. 6.**
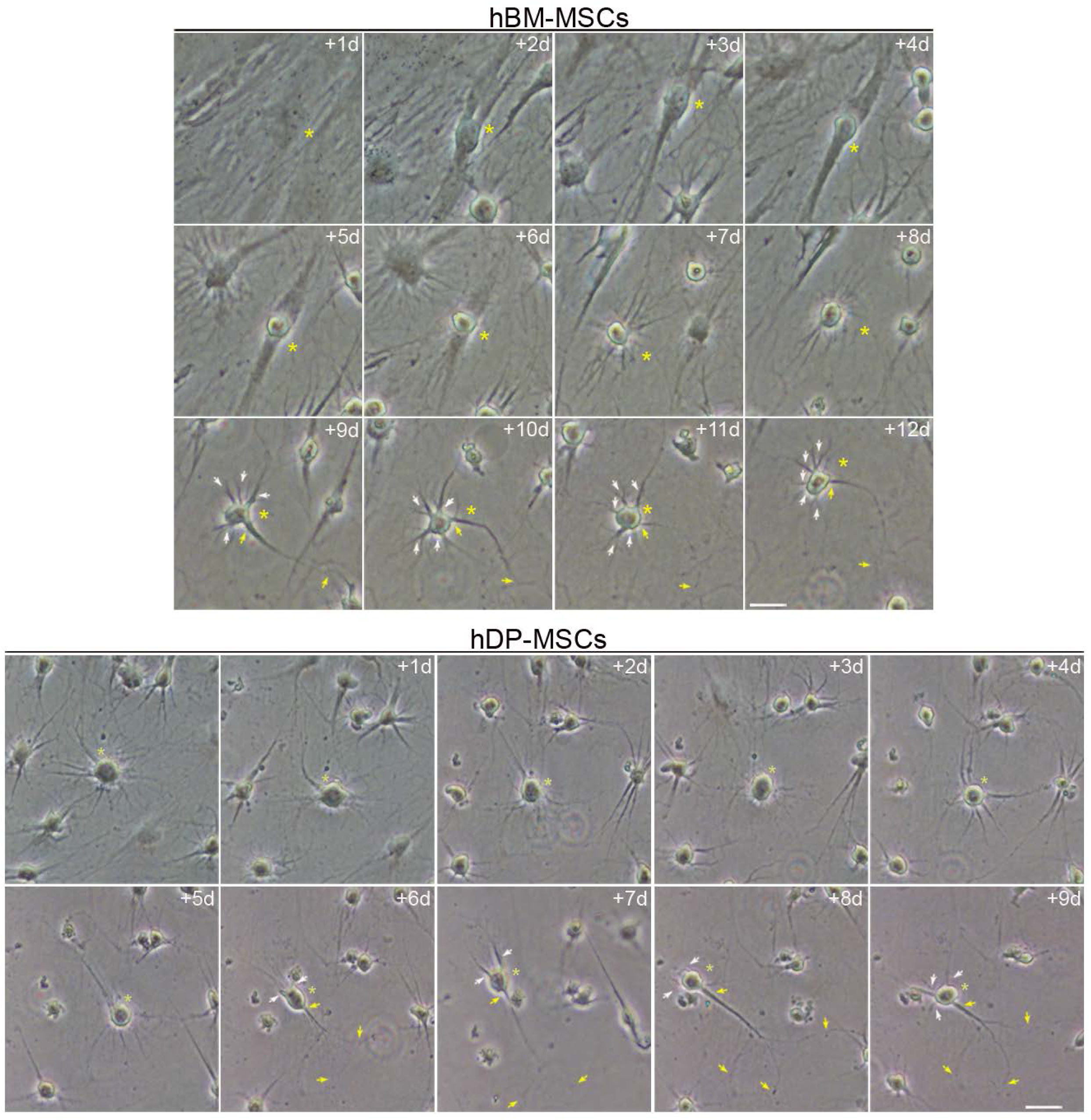
Neuron-like cells derived from human somatic cells exhibit neuronal polarization. Time-lapse imaging reveals the emergence of new neurite-like extensions sprouting directly from the cell bodies of these newly hBM-MSCs and hDP-MSCs derived-round cells. Subsequently, we observed the formation of cells that exhibit a long neurite-like extension (yellow arrows) and multiple short neurite-like extensions (white arrows) extending from the opposite side of the cell body, resulting in cells that resemble neuronal morphology. Scale bar: 10 μm. The number at the top indicates the time since the time-lapse image began. Elapsed time is displayed in days.

**Fig. 7.**
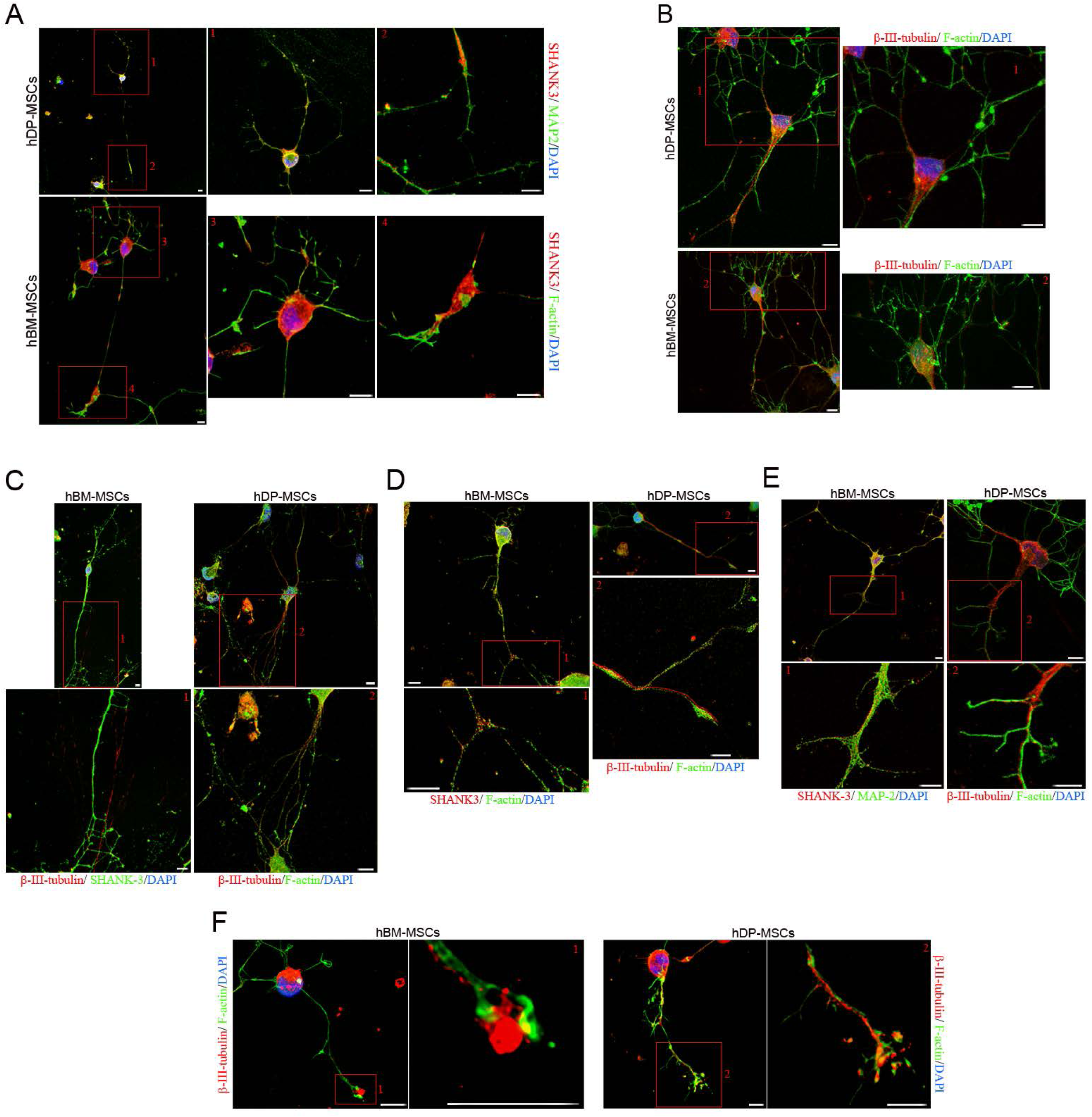
Neuron-like cells derived from hMSCs are highly polarized cells. (**A**) Analysis of the cytoskeleton revealed that neuron-like cells derived from hBM-MSCs and hDP-MSCs are highly polarized cells that develop distinctive extensions resembling dendrites and axons. (**B**) Cytoskeletal protein β-III tubulin and F-actin staining shown that neuron-like cells develop dendrite-like processes with a similar spatial organisation as that described in neurons. Neuron-like cells also display long neurite-like extension with various types of axonal branch-like structures that mimic neurons, including terminal arborization (**C**), bifurcation (**D**), and collateral formation (**E**). Cytoskeletal protein β-III tubulin and F-actin staining shown that neuron-like cells produce long neurite-like extension featuring growth cones at their tips, with a spatial organisation similar to that described in neurons. Scale bar: 5 μm.

**Fig. 8.**
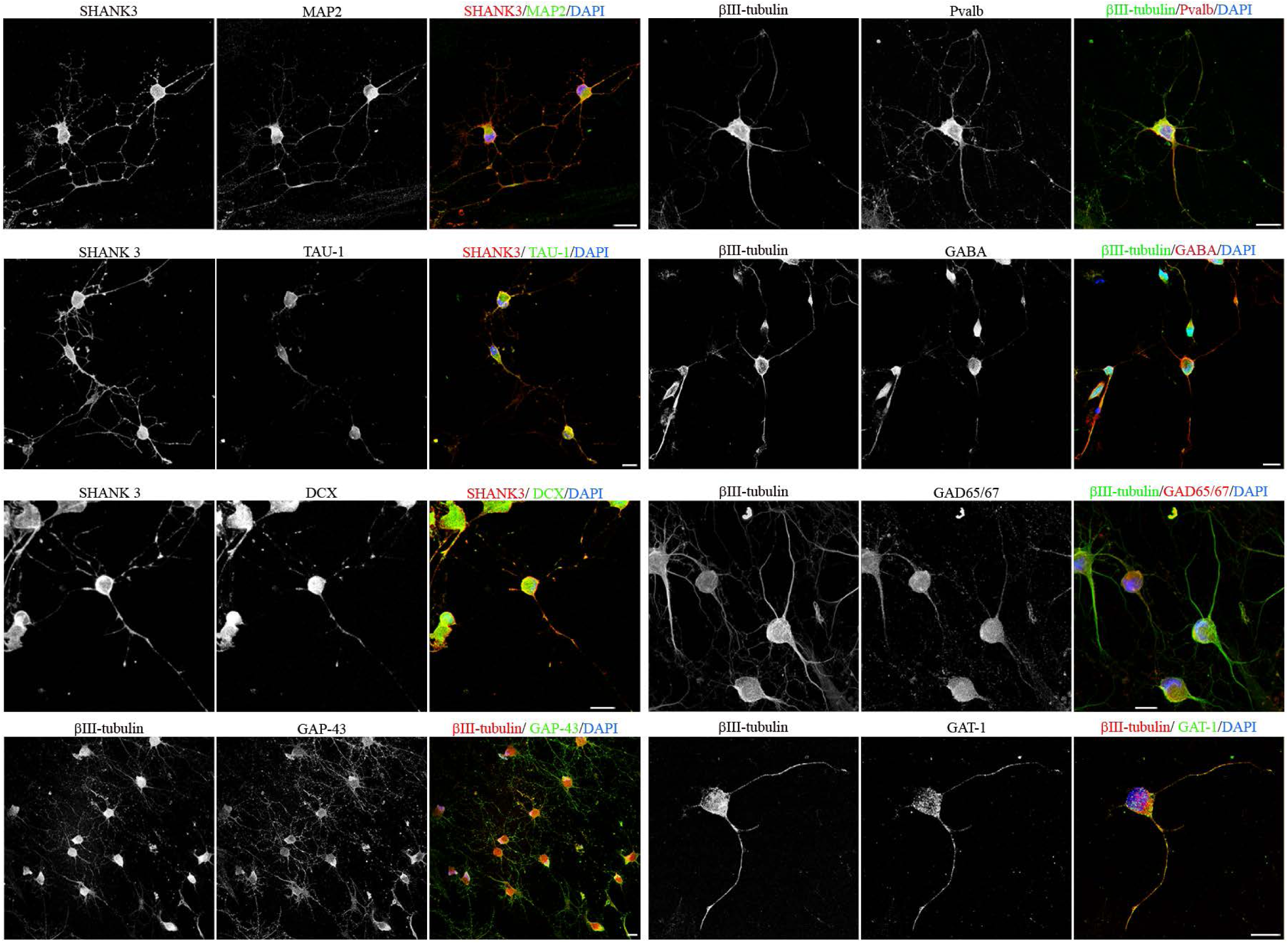
Neuron-like cells derived from hBM-MSCs express neuronal markers. Immunocytochemical analysis shown that neuron-like cells derived from hBM-MSCs express neuronal markers. Scale bar: 10 μm.

**Fig. 9.**
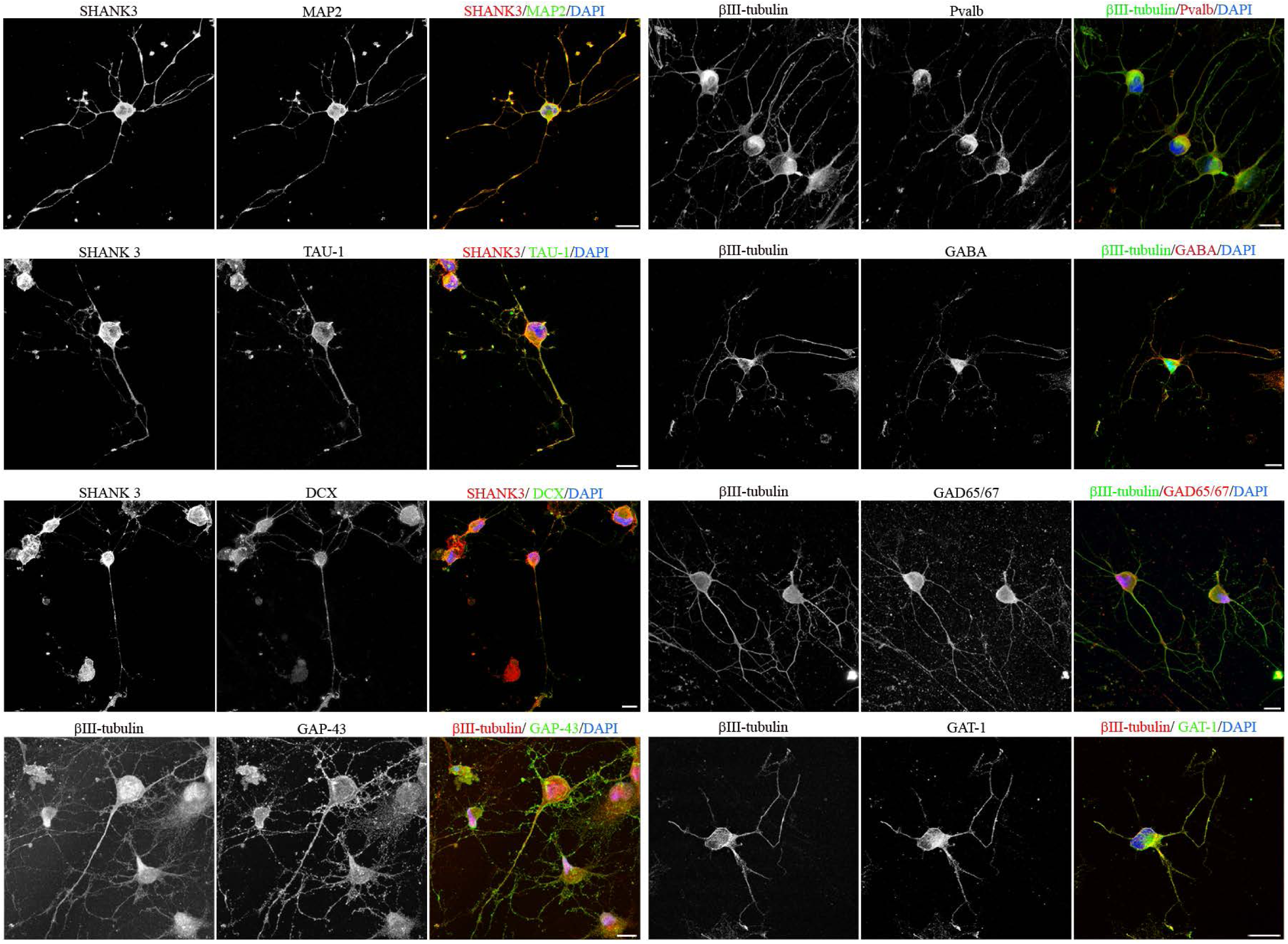
Neuron-like cells derived from hDP-MSCs express neuronal markers. Immunocytochemical analysis shown that neuron-like cells derived from hDP-MSCs express neuronal markers. Scale bar: 10 μm.

**Fig. 10.**
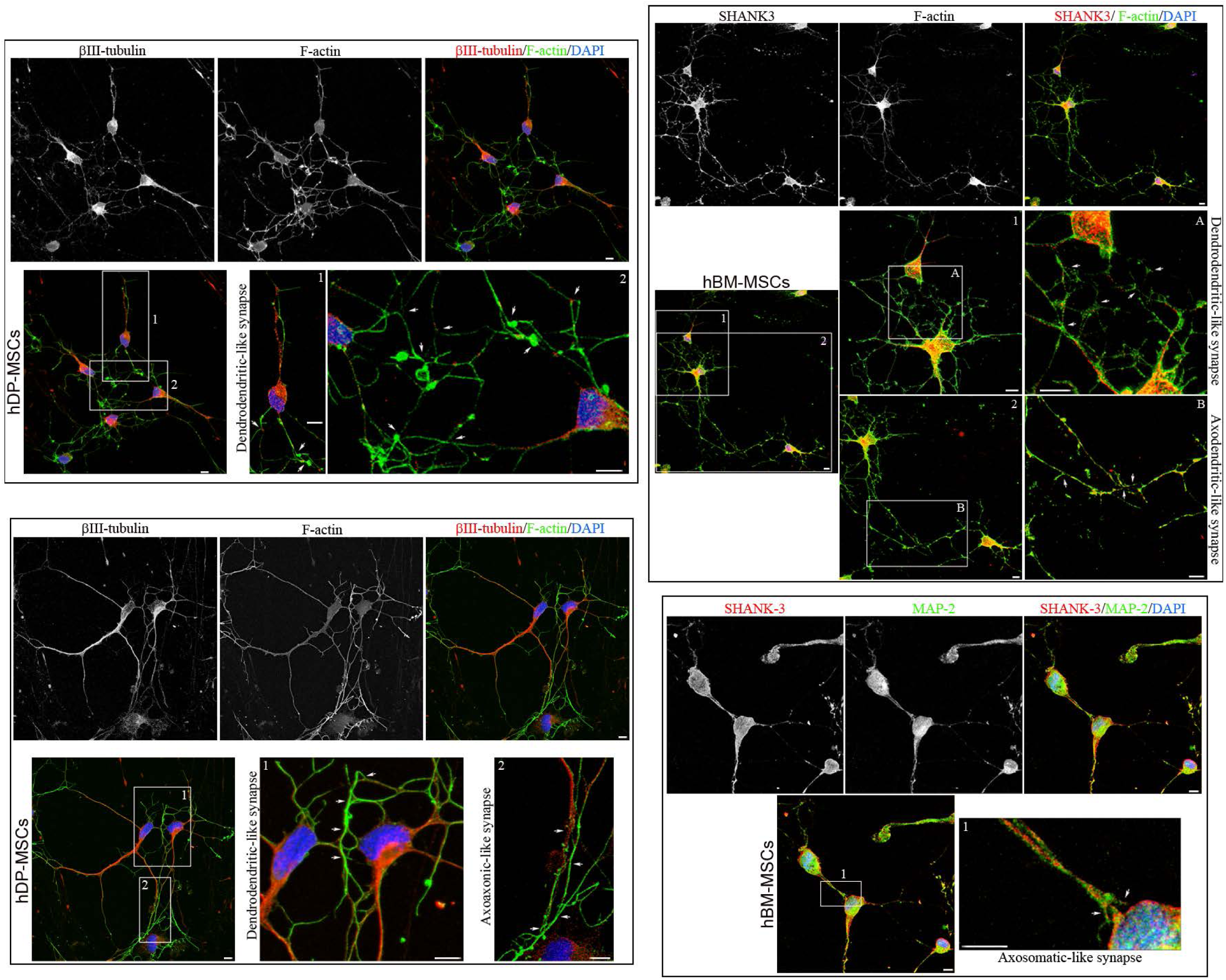
Neuron-like cells derived from hMSCs established connections with one another forming interconnected networks. Immunocytochemical analysis shown that neuron-like cells derived from hBM-MSCs and hDP-MSCs were interconnected via various types of connections, including dendrodendritic, axodendritic, axosomatic and axoaxonic-like contacts. The white arrows show the connection points between these cells. Scale bar: 5 μm.

**Fig. 11.**
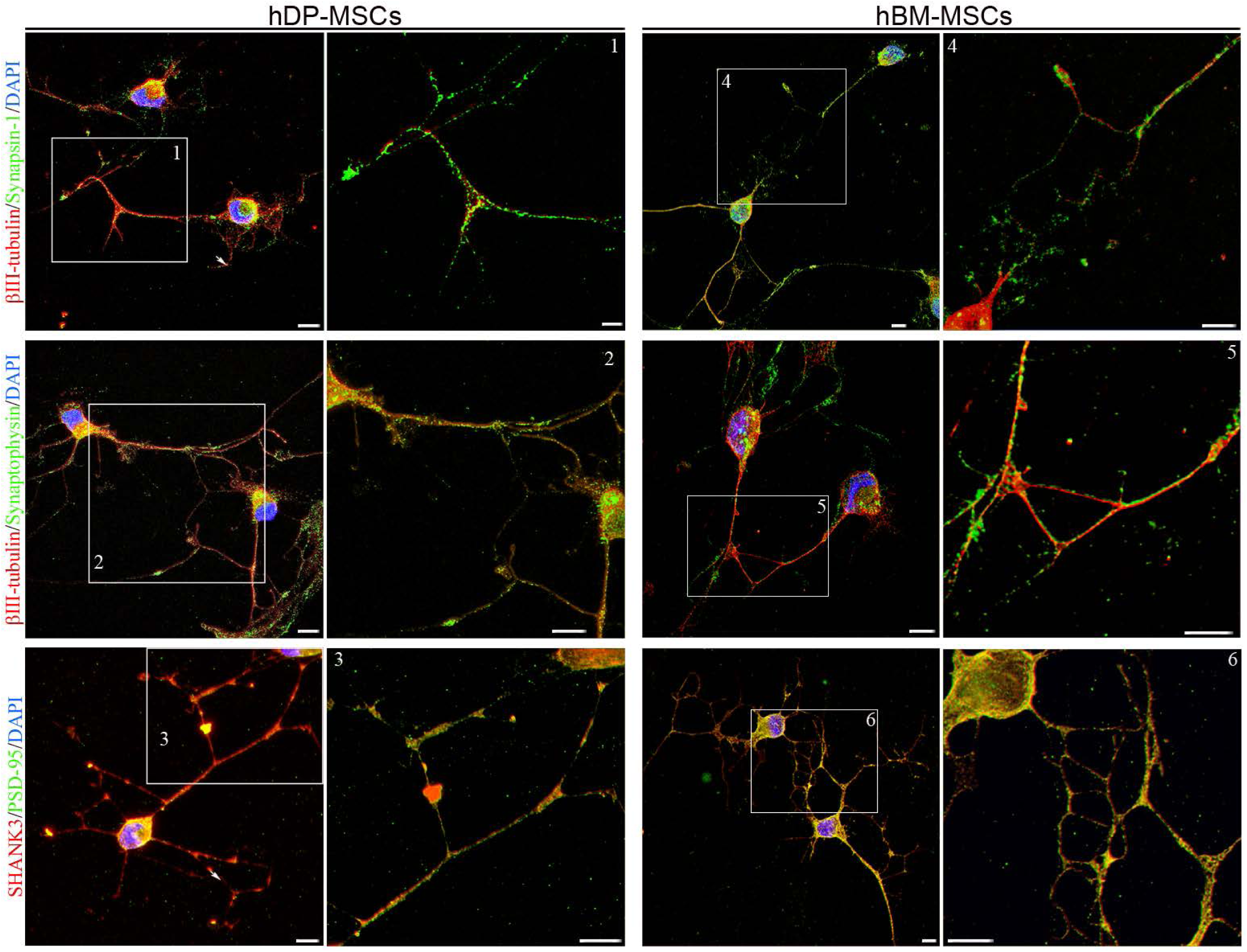
Neuron-like cells derived from hMSCs express synapse-associated proteins. Immunocytochemical analysis shown that neuron-like cells derived from hBM-MSCs and hDP-MSCs express synapse-associated proteins. Scale bar: 5 μm.

**Fig. 12.**
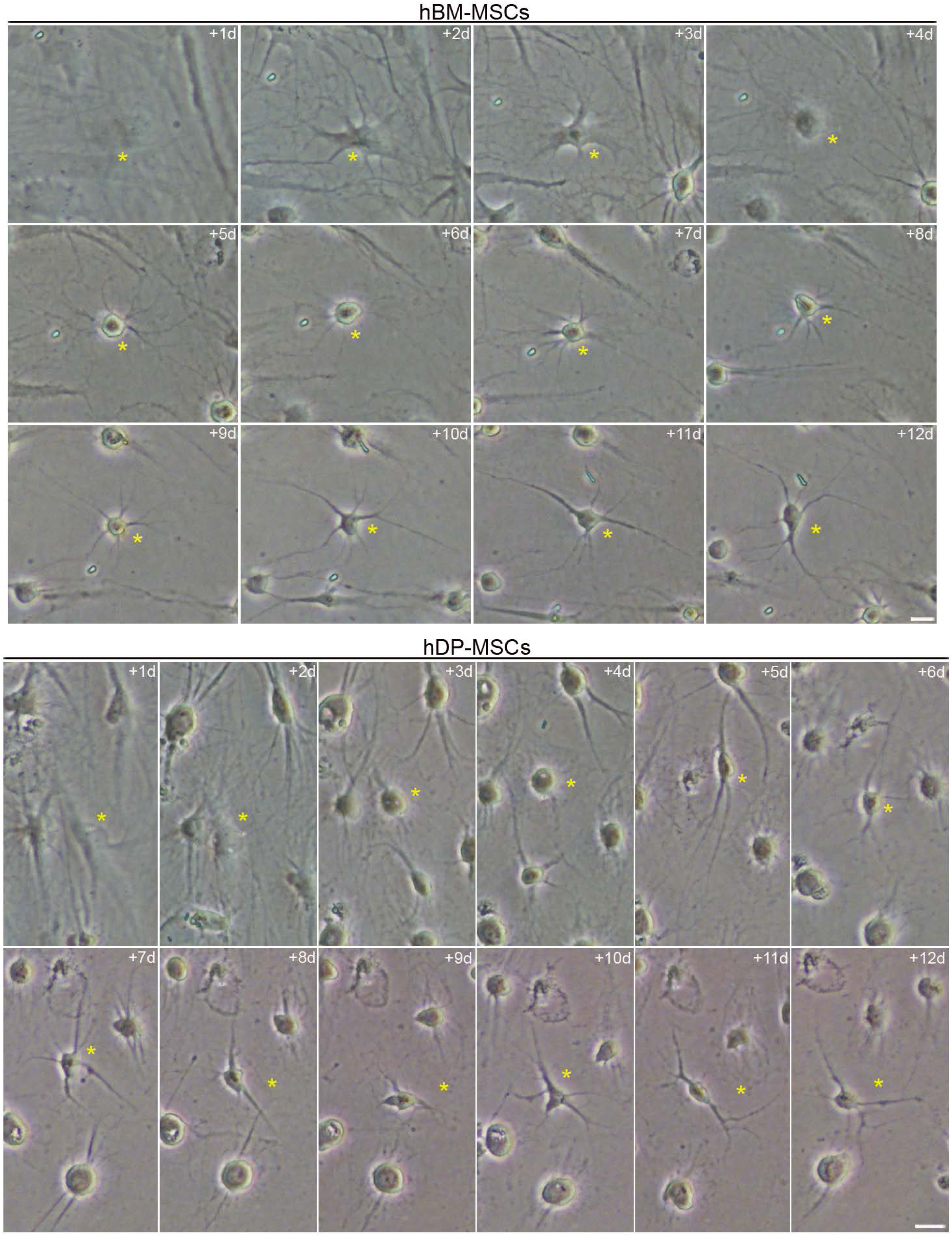
Neuron-like cells derived from hMSCs exhibit plasticity. Time-lapse imaging shown that neuron-like cells derived from hBM-MSCs and hDP-MSCs undergo progressive processes of neurite remodelling involving physical changes in their architecture. Scale bar: 10 μm. The number at the top indicates the time since the time-lapse image began. Elapsed time is displayed in days.

**Fig. 13.**
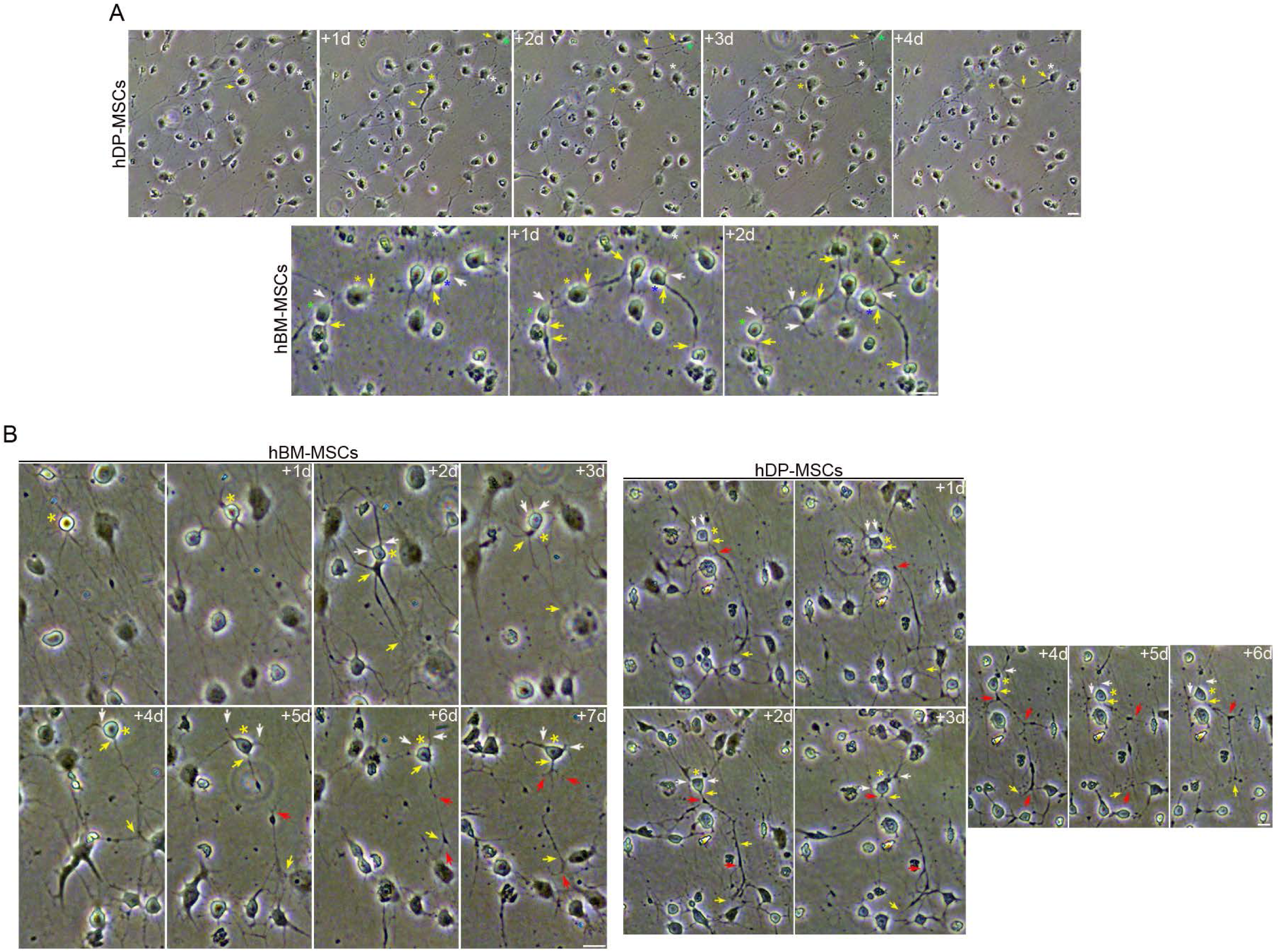
Neuron-like cells derived from hMSCs exhibit structural plasticity. Time-lapse imaging shown that neuron-like cells derived from hBM-MSCs and hDP-MSCs are capable of physically remodelling their connections through the growth, retraction (**A**) and branching of axon-like processes (**B**) and dendrite-like processes (**A**, **B**) to establish new connections or reorganize existing ones. The white and yellow arrows indicate, respectively, dendrite-like processes and axon-like processes. The red arrows indicate branching points of axon-like processes. Scale bar: 10 μm. The number at the top indicates the time since the time-lapse image began. Elapsed time is displayed in days.

**Fig. 14.**
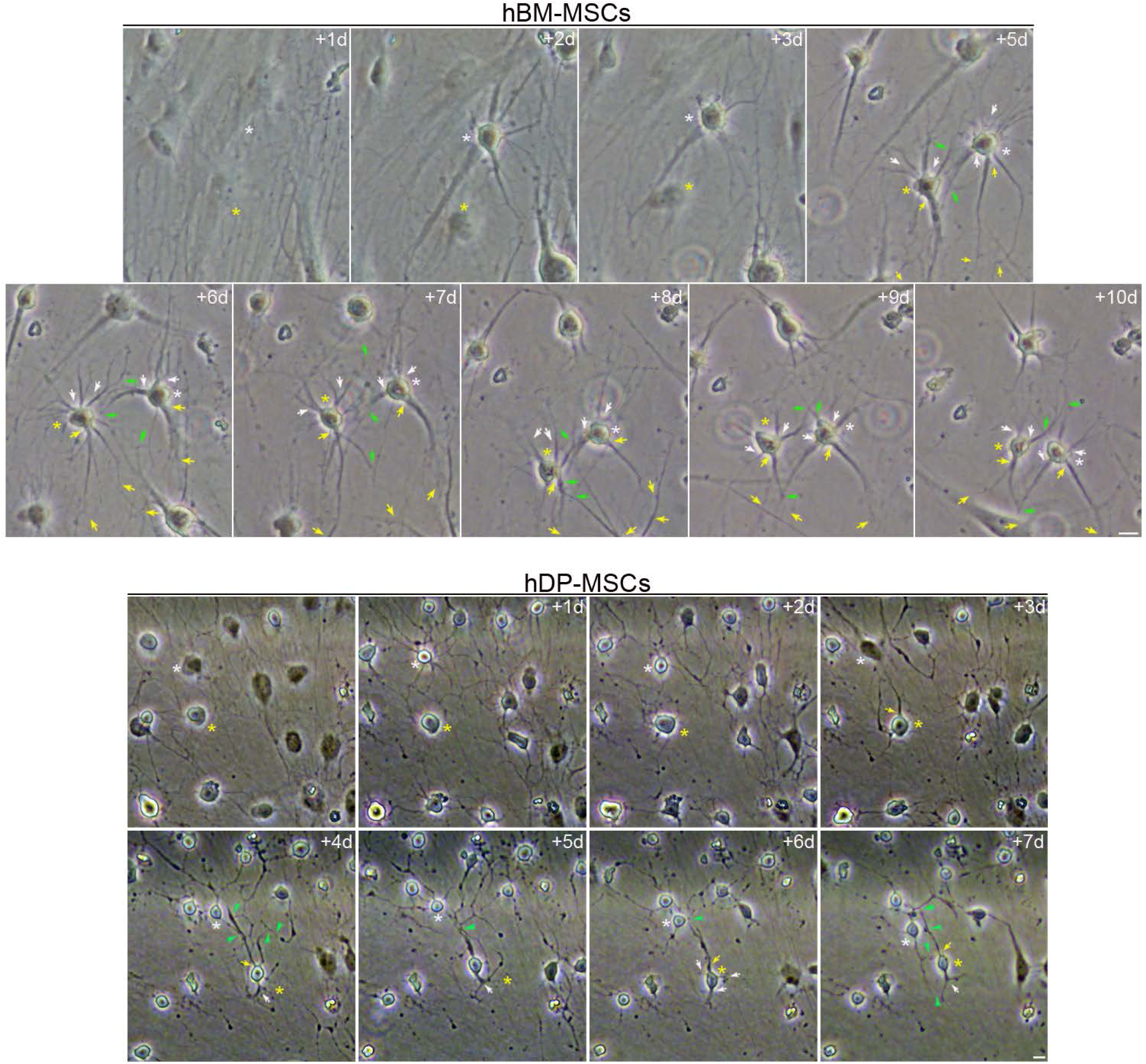
Neuron-like cells derived from hMSCs display neurite-like interactions that exhibit connection plasticity. Time-lapse imaging indicate that neuron-like cells derived from hBM-MSCs and hDP-MSCs develop distinctive extensions resembling dendrites (white arrows) and axons (yellow arrows) that form plastic, changing connections. The green arrows indicate the connection points between these cells. Scale bar: 10 μm. The number at the top indicates the time since the time-lapse image began. Elapsed time is displayed in days.

In cultured primary neurons, the formation of interconnected networks is the functional endpoint of neuronal differentiation [35, 36]. This process transforms isolated cells into a synchronized, communicating circuit. As mentioned above, time-lapse monitoring and morphological analysis shown that neuron-like cells derived from hBM-MSCs and hDP-MSCs establish interconnected networks (Figs. 1, 2, 3, 4, 5). Immunocytochemical analysis also revealed that neuron-like cells were interconnected via various types of connections, including dendrodendritic, axodendritic, axosomatic and axoaxonic-like contacts (Fig. 10). Synapse-associated proteins, such as Synaptophysin, Synapsin1, PSD-95 and SHANK-3 are found in neuron-like cells derived from hBM-MSCs and hDP-MSCs (Fig. 11).

Neurons exhibit plasticity, which means that they can reorganise their structure, function and connections; this involves the growth, retraction and branching of dendrites and axons to establish new connections or reorganize existing ones [40]. These mechanisms allow neurons to establish functional networks during development and maintaining brain function, including learning and memory, as well as in recovery from brain injury [41, 42].

Time-lapse imaging reveals that neuron-like cells derived from hBM-MSCs and hDP-MSCs exhibit remarkable morphological plasticity as they undergo progressive neuritic remodeling processes that involve physical changes in their neuron-like architecture (Fig.12). Time-lapse imaging also shown that neuron-like cells possess the capacity for structural plasticity, allowing them to physically remodel their connections (Fig.13). This structural plasticity involves the growth (Fig. 13A), retraction (Fig. 13A) and branching of axon-like processes (Fig. 13B) and dendrite-like processes (Fig. 13) to establish new connections or reorganize existing ones. Observations from time-lapse microscopy also indicate that neuron-like cells connect through various neurite-like interactions that exhibit connection plasticity (Fig.14).

Action potentials are regarded as one of the most important functional markers of neuronal identity in neurons derived from human somatic cells, as they represent the culmination of a successful reprogramming or differentiation process. Unfortunately, in this study we have not had the opportunity to perform patch-clamp electrophysiological techniques or to use multi-electrode arrays to demonstrate that neuron-like cells derived from hBM-MSCs and hDP-MSCs display functional excitability. However, a recent study has shown that hDP-MSCs can be differentiated into neuron-like cells that exhibit functional excitability, as demonstrated by patch-clamp experiments [43]. Further research is needed to demonstrate that hBM-MSCs display functional excitability.

Our results showed that neuron-like cells derived from hMSCs exhibit neuronal polarization, express specific neuronal and synaptic markers, form interconnected networks and exhibit structural plasticity. These findings confirm our previous results [26–29] and provide further evidence that hMSCs can be directly converted into cells with neuronal features.

### The differentiation of human somatic cells into neuron**-**like cells involves a transition through a transient intermediate state

In previous publications [29, 30], we showed that the conversion of hBM-MSCs into neuron-like cells involves a transition through a transient intermediate state. During this cellular state, the cells adopt a spherical morphology, and the nuclei are highly dynamic: they move through the cytoplasm, change shape and position, and even form cellular protrusions as they attempt to make contact with surrounding cells, likely to sense their microenvironment. We also observed that intermediate cells can preserve their spherical shape for several days without assuming new fates, or they can adopt alternative fates; they gradually adopted a neural-like morphology through active neurite extension or re-differentiated back to the mesenchymal fate. Furthermore, we found that intermediate cells undergo rapid and repeated lineages transitions without cell division [29].

To better understand whether these cells truly represent an intermediate stage prior to neuronal polarization, we investigated whether nuclear movement is observed exclusively in these newly formed round cells or whether it also occurs during neuronal polarization. Time-lapse images revealed the presence of round cells derived from hBM-MSCs and hDP-MSCs, in which cellular protrusions caused by nuclear movement were observed (Fig. 15A). However, no transient cellular protrusions were observed, as the round cells gradually acquired a neuron-like morphology (Fig. 15B). Although further studies using live-cell nuclear fluorescence labelling and time-lapse microscopy are needed to determine whether the cell nucleus actually ceases to move during neural polarization, these results suggest that these rounded cells, in which the cell nucleus is highly dynamic, represent an intermediate stage of differentiation prior to neuronal polarization.

**Fig. 15.**
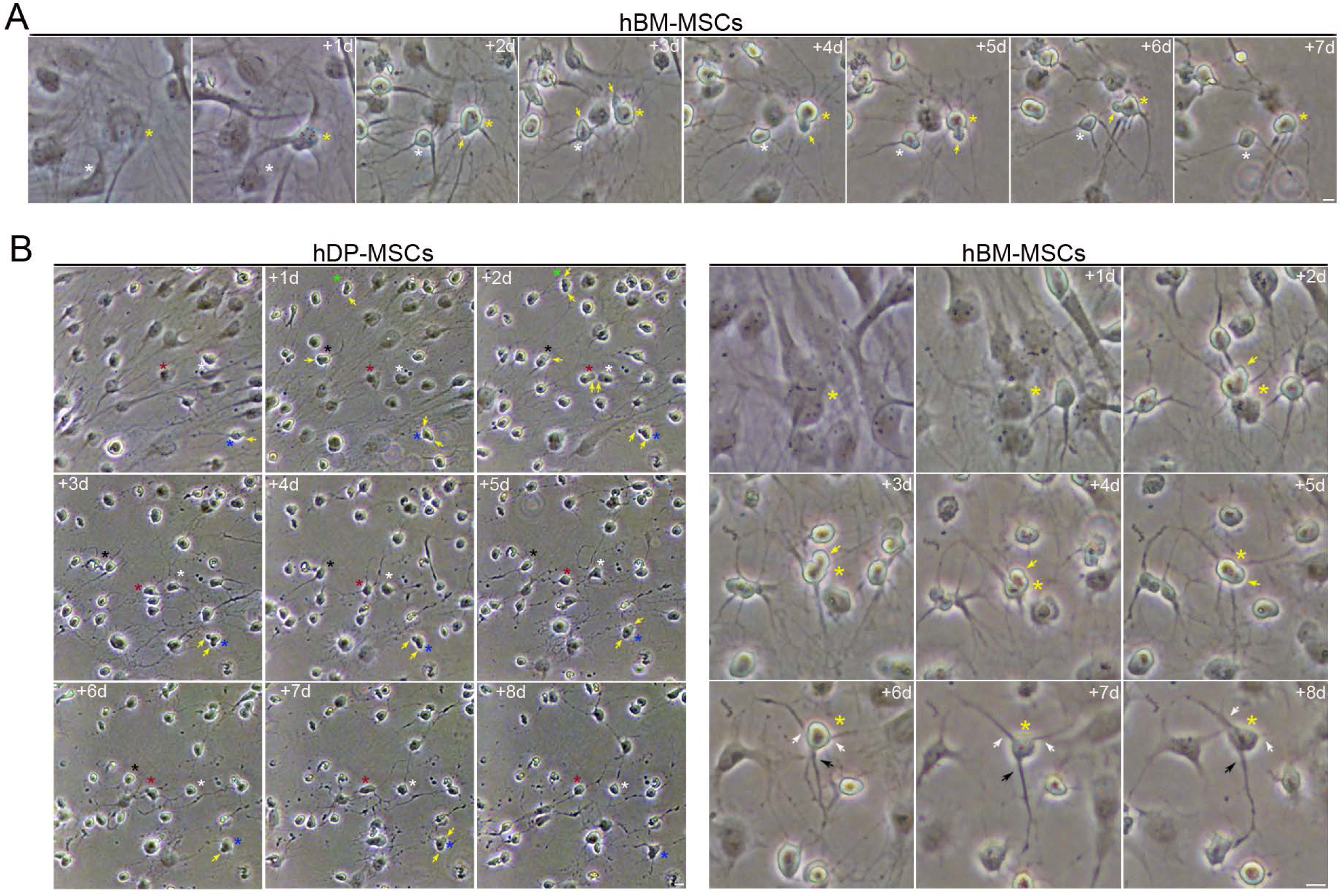
Newly formed round cells represent an intermediate stage of differentiation prior to neuronal polarization. Time-lapse images revealed the presence of newly formed round cells derived from hMSCs, in which cellular protrusions caused by nuclear movement were observed (**A**). However, no transient cellular protrusions were observed, as the round cells gradually acquired a neuron-like morphology (**B**). The yellow arrows indicate cellular protrusions caused by nuclear movement. The white and black arrows indicate, respectively, dendrite-like processes axon-like processes. Scale bar: 10 μm. The number at the top indicates the time since the time-lapse image began. Elapsed time is displayed in days.

Previous studies have reported that direct reprogramming (or transdifferentiation) of somatic cells into neurons bypasses the traditional stages of pluripotency or the intermediate stages of development [9–19, 23, 24]. MEFs and HDFa are the most used starting cells for direct reprogramming to generate functional neurons. However, it is important to note that these reports [9–19] do not include time-lapse experiments, which allow for direct and continuous visualization of cellular fate conversion, thereby overcoming the limitations of fixed-cell analysis, which often overlooks short-lived transient states.

In this study, we also examined whether the direct reprogramming of HDFa into neurons involves passing through a transient intermediate state. Time-lapse images revealed that, when HDFa were exposed to a neural induction medium, they exhibited sequences of neuronal morphological development highly similar to those of hBM-MSCs and hDP-MSCs (Fig. 16). Time-lapse microscopy also show that it is possible to observe HDFa that reshape from a flat to a spherical morphology. Subsequently, we noted the formation of cells that exhibit a long neurite-like extension and multiple short neurite-like extensions extending from the opposite side of the cell body, resulting in cells that resemble neuronal morphology (Fig. 17). It is important to note that no cell division is observed during the conversion HDFa into neuron-like cells (Figs. 17-20).

**Fig. 16.**
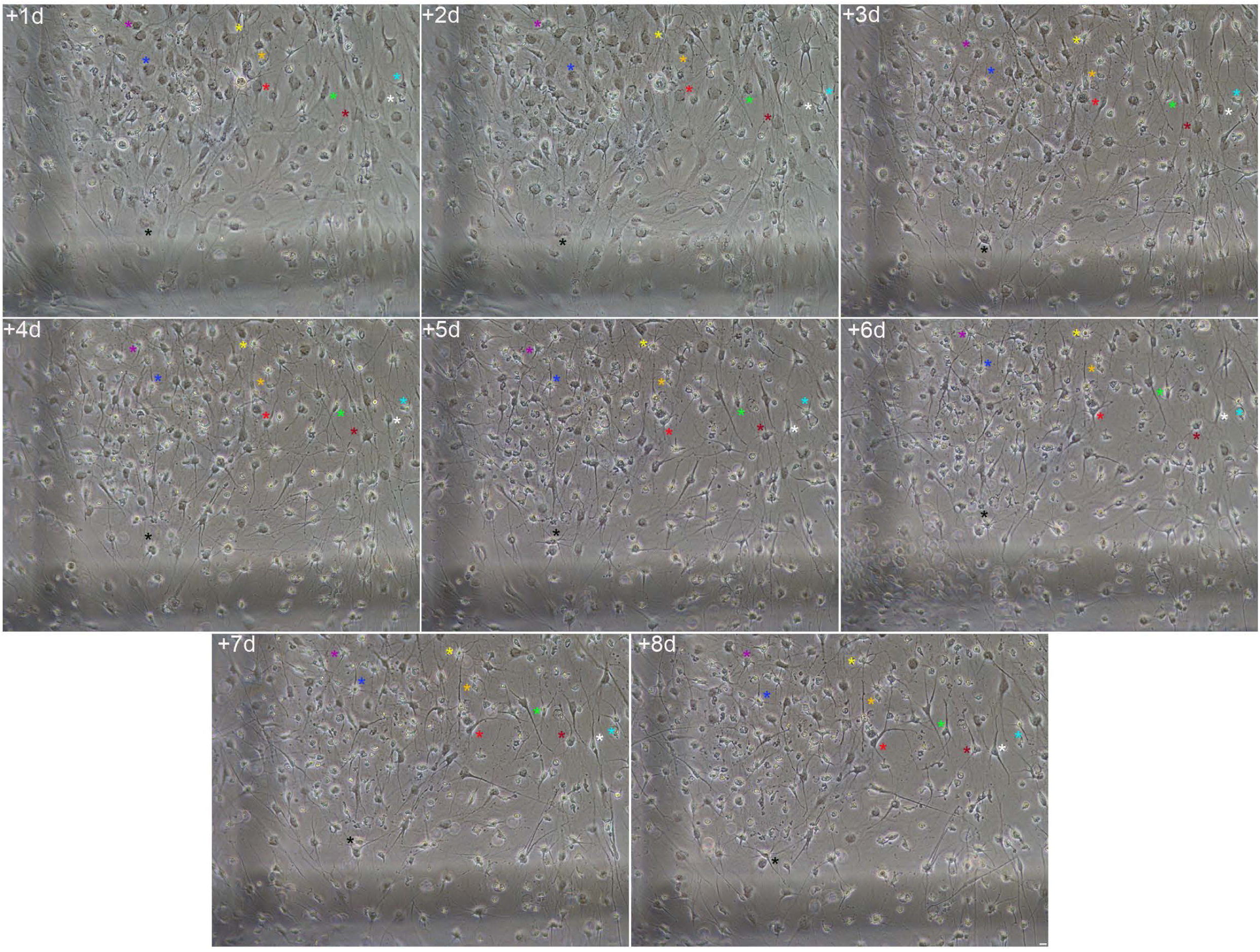
Chemical conversion of HDFa to neuron-like cells. Time-lapse imaging revealed that when adult HDFa were exposed to neural induction medium, they reshaped from a flat to a spherical morphology. We then observed the emergence of new neurite-like extensions sprouting directly from the cell bodies of these rounded cells. Subsequently, these cells gradually acquired a neuron-like morphology and established connections with one another through various types of interactions. Scale bar: 10 μm. The number at the top indicates the time since the time-lapse image began. Elapsed time is displayed in days.

**Fig. 17.**
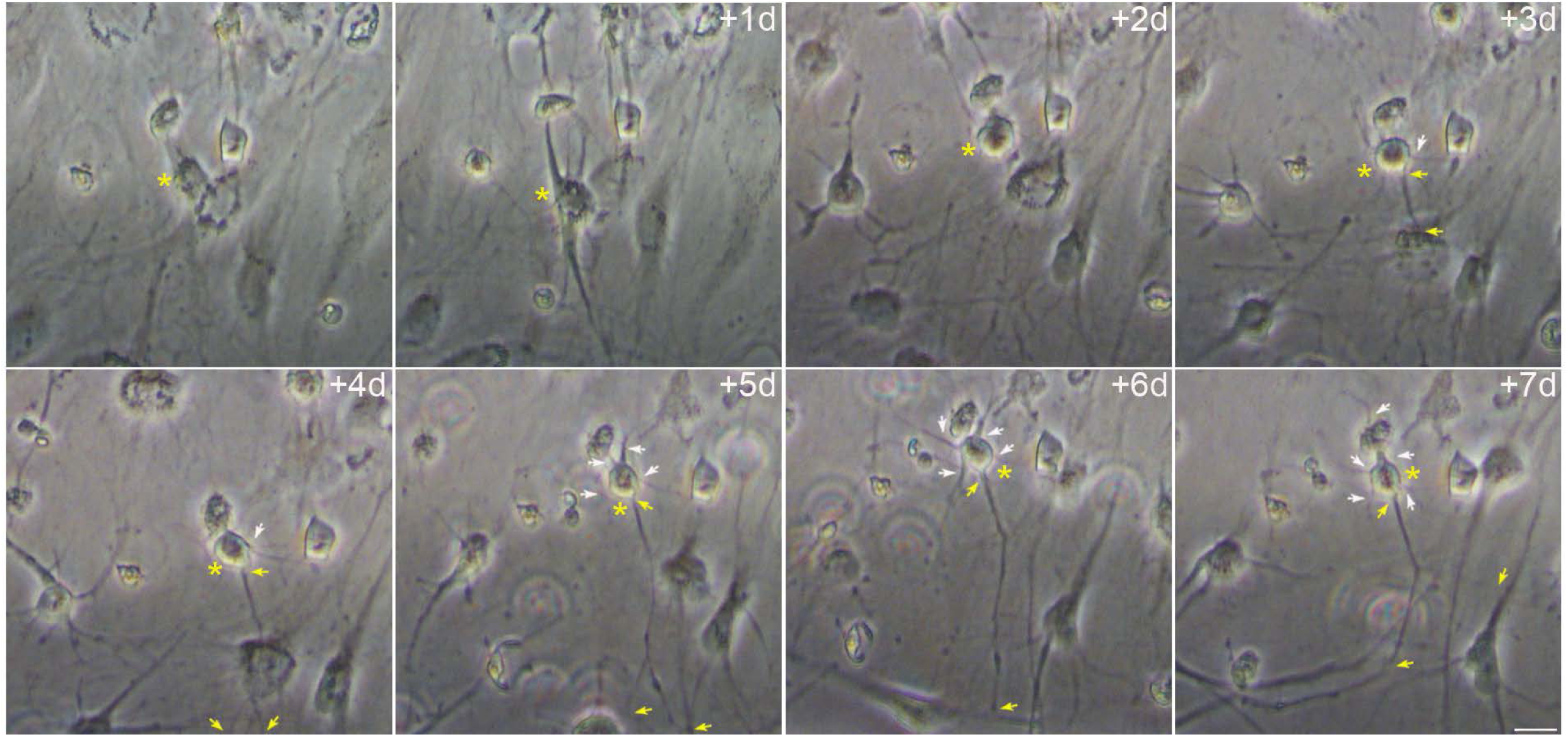
Neuron-like cells derived from HDFa exhibit neuronal polarization. Time-lapse imaging reveals the emergence of new neurite-like extensions sprouting directly from the cell bodies of these newly formed round cells derived from HDFa. Subsequently, we observed the formation of cells that exhibit a long neurite-like extension (yellow arrows) and multiple short neurite-like extensions (white arrows) extending from the opposite side of the cell body, resulting in cells that resemble neuronal morphology. Scale bar: 10 μm. The number at the top indicates the time since the time-lapse image began. Elapsed time is displayed in days.

**Fig. 18.**
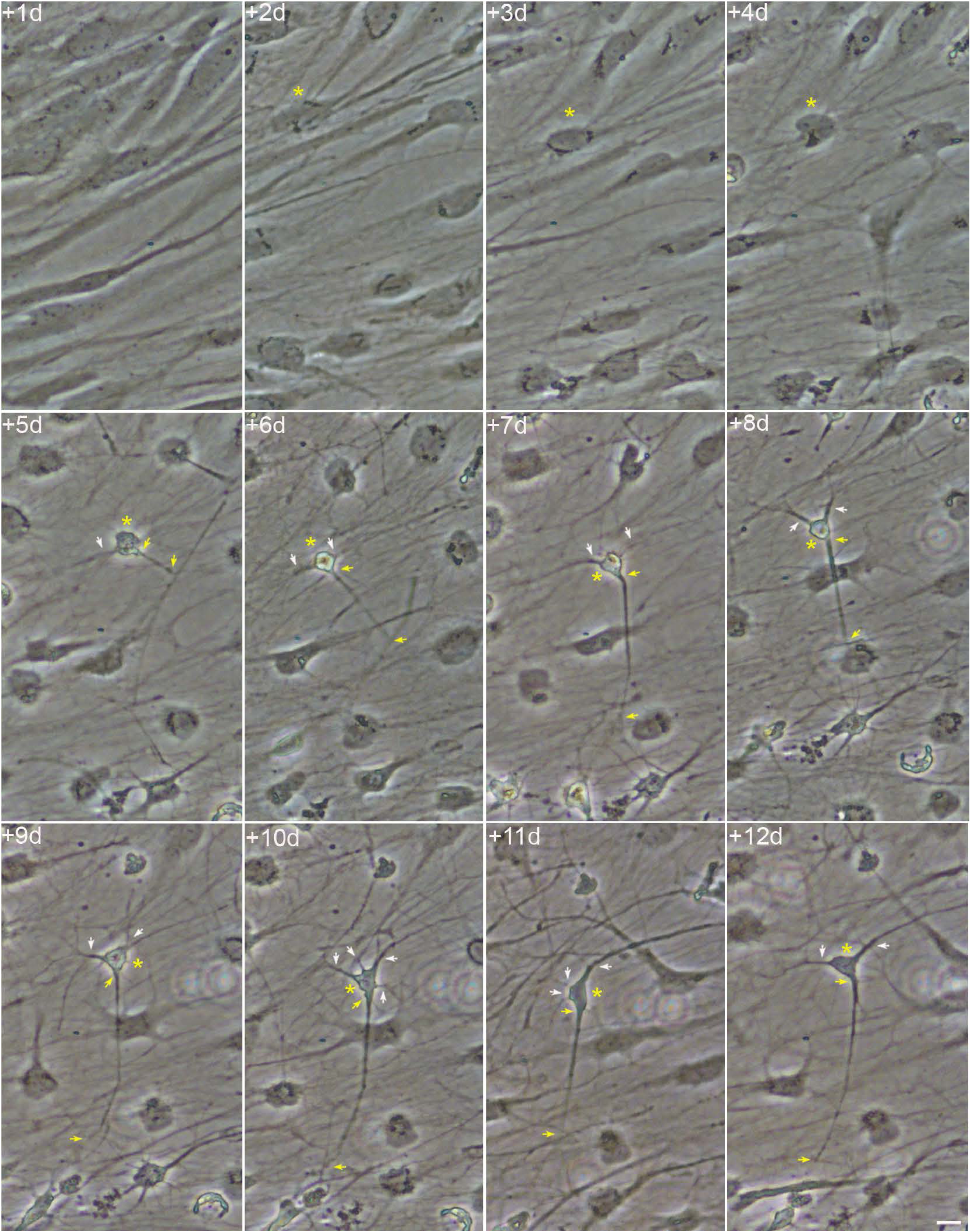
Neuron-like cells derived from HDFa exhibit morphological plasticity. Time-lapse imaging reveals that neuron-like cells derived from HDFa exhibit morphological plasticity as they undergo progressive neuritic remodelling processes that involve physical changes in their dendrite-like processes. The white and yellow arrows indicate, respectively, dendrite-like processes and axon-like processes. Scale bar: 10 μm. The number at the top indicates the time since the time-lapse image began. Elapsed time is displayed in days.

**Fig. 19.**
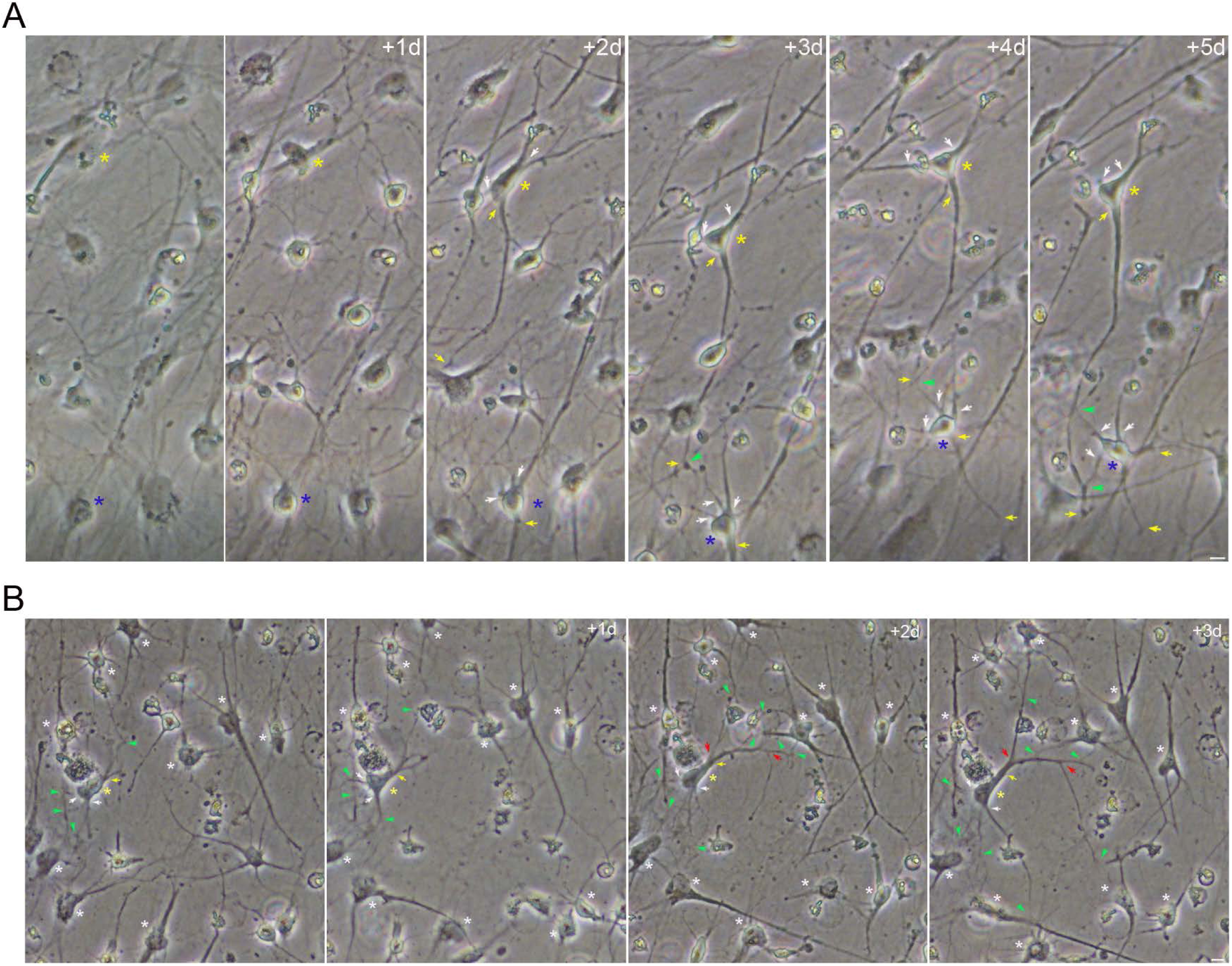
Neuron-like cells derived from HDFa display neurite-like interactions that exhibit connection plasticity. Time-lapse imaging indicate that neuron-like cells derived from HDFa develop distinctive extensions resembling dendrites (white arrows) and axons (yellow arrows) that form plastic, changing connections. The red arrows indicate branching points of axon-like processes. The green arrows indicate the connection points between cells. Scale bar: 10 μm. The number at the top indicates the time since the time-lapse image began. Elapsed time is displayed in days.

**Fig. 20.**
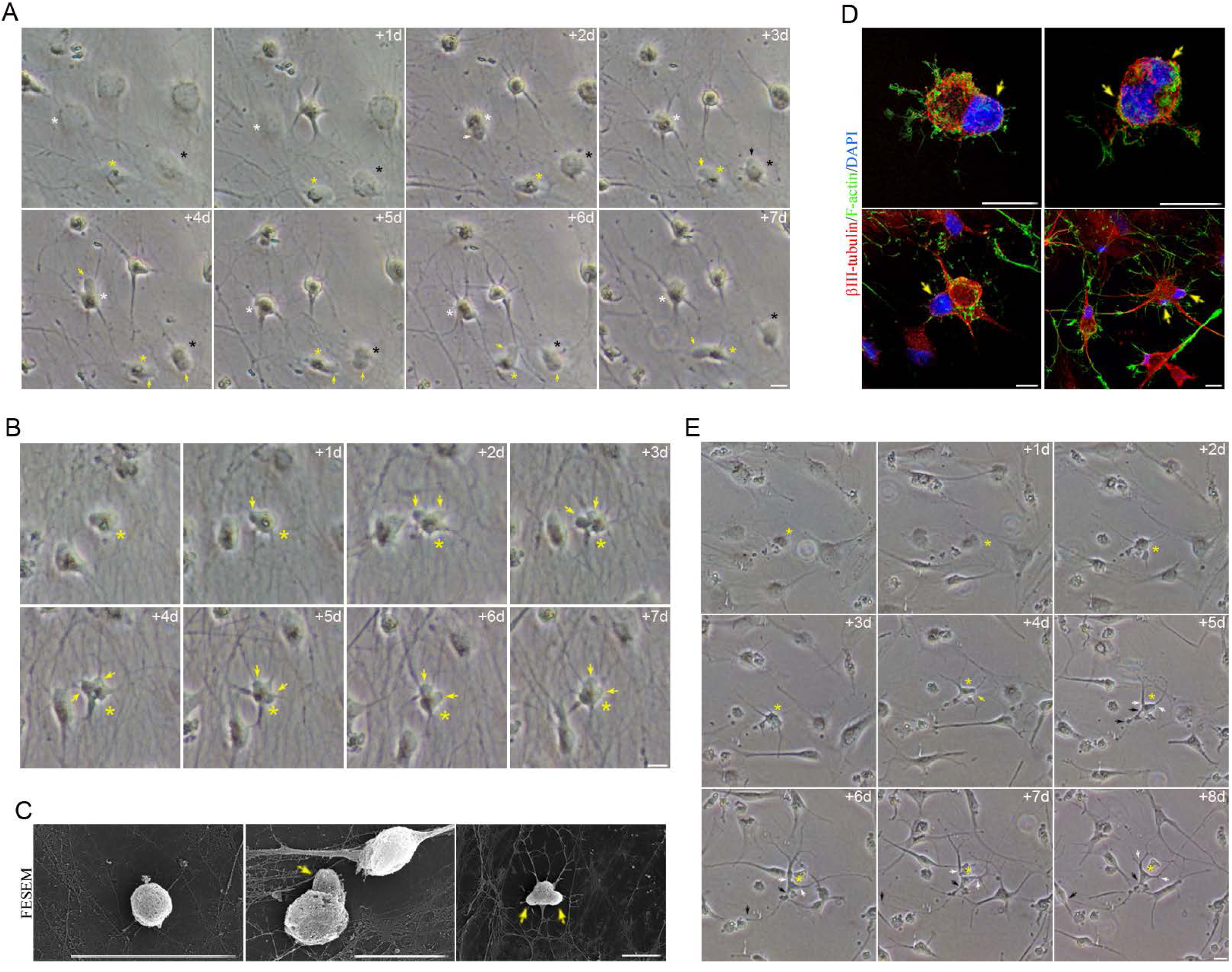
The chemical conversion of HDFa to neuron-like cells involves passing through a transient intermediate state. Time-lapse images revealed the presence of newly formed round cells derived from HDFa, in which one (**A**) or two (**B**) cellular protrusions caused by nuclear movement can be observed. (C) Scanning electron micrographs showed newly formed round cells derived from HDFa, in which nuclear movements generate only one cell protrusion or two cellular protrusions. (**D**) Immunocytochemical analyses confirm the presence of newly formed round cells with non-lobulated nuclei and lobulated nuclei that form cellular protrusions. (**E**) Time-lapse imaging also indicate that no transient cellular protrusions were observed, as the round cells gradually acquired a neuron-like morphology. The yellow arrows indicate cellular protrusions caused by nuclear movement. The white and black arrows indicate, respectively, dendrite-like processes and axon-like processes. Scale bar: 10 μm. The number at the top indicates the time since the time-lapse image began. Elapsed time is displayed in days.

Time-lapse imaging also reveals that neuron-like cells derived from HDFa exhibit morphological plasticity as they undergo progressive neuritic remodelling processes that involve physical changes in their dendrite-like processes (Fig. 18). Observations from time-lapse microscopy indicate that neuron-like cells connect through various neurite-like interactions that exhibit connection plasticity (Fig. 19). This structural plasticity involves the growth (Fig. 19A) and branching of axon-like processes (Fig. 19B), as well as dendrite-like processes (Fig. 19) to establish new connections or reorganize existing ones.

Time-lapse images also revealed the presence of newly formed round cells derived from HDFa, in which cellular protrusions caused by nuclear movement were observed (Fig. 20). We noted cells in which nuclear movements generate only one cell protrusion (Fig. 20A) or two cellular protrusions (Fig. 20B). In a previous publication [30], we have shown that when an hBM-MSC-derived intermediate cell has a nucleus without lobes, its movement within the cell generates only one cell protrusion. However, if the hBM-MSC-derived intermediate cell has a lobed nucleus, it will generate one or two cellular protrusions depending on how it moves within the cell. Scanning electron micrographs also showed newly formed round cells derived from HDFa, in which nuclear movements generate only one cell protrusion or two cellular protrusions (Fig. 20C). Immunocytochemical analyses confirm the presence of newly formed round cells with non-lobulated nuclei and lobulated nuclei that form cellular protrusions (Fig. 20D). Observations from time-lapse microscopy also indicate that no transient cellular protrusions were observed, as the round cells gradually acquired a neural-like morphology (Fig. 20E).

Our results showed that when somatic cells isolated from various sources in the human body, are exposed to the neural induction medium, they change shape and transition to an intermediate cellular state before adopting neural-like morphologies. During this cellular state, the cells adopt a spherical morphology, and the nuclei are highly dynamic: they move through the cytoplasm, change shape and position, and even form cellular protrusions as they attempt to make contact with surrounding cells, likely because they are sensing their environment. These findings confirm our previous results [27–30] and provide further evidence that the conversion of human somatic cells into neurons involves passing through a transient intermediate state.

It is important to note that, in our opinion, cells with lobulated or binucleated nuclei may also appear to be present in direct reprogramming experiments conducted using both the overexpression of transcription factors [9] (their Fig. S8B) as well as small-molecule-based protocols [18] (their Figs. 1D, S1D, S2A, S2B).

## Discussion

Traditionally, lineage commitment and differentiation have been considered to be unidirectional and irreversible [44]. However, landmark experiments such as somatic cell nuclear transfer into enucleated oocytes and cell fusion of pluripotent cells with differentiated cells demonstrated that differentiated cells are not irreversibly committed to their fate [45]. More recent work has built on these findings and has shown that the overexpression of transcription factors or non-coding RNAs, as well as the use of small molecules, can directly induce one specific cell type from another, even between distantly related as cells from different germ layers [5–19, 45–47]. These findings challenge the principles of irreversibility and restriction of the germ layers [24, 48], as well as the classical definitions of stem cells [49] and pluripotency [24]. Although cell lineage conversion can be experimentally induced using cell reprogramming and transdifferentiation techniques, it is important to note that changes involving a shift in cell fate, such as dedifferentiation, transdetermination and transdifferentiation, are also observed in physiological processes that occur during tissue regeneration and repair following injury [50], as well as in conditions such cancer, where they contribute to tumor progression and metastasis [51]. Therefore, future research on the conversion and reprogramming of cell lineages is not only essential for understanding developmental processes and cell fate decisions, but also, for translating these insights into regenerative medicine applications.

Our current research aims to improve our understanding of direct reprogramming (or transdifferentiation) of human somatic cells into neuronal cells using small molecule-based protocols. In previous publications, we showed that the conversion of hBM-MSCs into neuron-like cells involves a transition through a transient intermediate state [29, 30]. During this cellular state, the cells adopt a spherical morphology, and the nuclei are highly dynamic: they move through the cytoplasm, change shape and position, and even form cellular protrusions as they attempt to make contact with surrounding cells, likely to sense their microenvironment. We also observed that intermediate cells can preserve their spherical shape for several days without assuming new fates, or they can adopt alternative fates; they gradually acquired a neural-like morphology through active neurite extension or re-differentiated back to the mesenchymal state. Furthermore, we found that intermediate cells undergo rapid and repeated lineages transitions without cell division [29]. Therefore, these intermediate cells, which differ from both the original mesenchymal cells and the final neural like-cells, may represent a previously underappreciated transitional intermediate stage in the transdifferentiation process. Previous studies reported that direct reprogramming of somatic cells into neurons bypasses the traditional stages of pluripotency or the intermediate stages of development [9–19, 23, 24]. MEFs and HDFa are the most used starting cells for direct reprogramming to generate functional neurons. However, it is important to note that these reports [9–19] do not include time-lapse experiments, which allow for continuous visualization of cellular fate conversion, thereby overcoming the limitations of fixed-cell analysis, which often overlooks short-lived transient states.

In this study, we conducted long-term live-cell imaging to observe the conversion of hBM-MSCs, hDP-MSCs and HDFa into neuron-like cells. Time-lapse analysis allows us not only to determine whether the direct reprogramming of somatic cells into neurons actually involves passing through a transient intermediate state, but also to provide additional evidence supporting the authenticity of human-derived neurons generated from hMSCs. Our results showed that when hBM-MSCs, hDP-MSCs and HDFa are exposed to the neural induction medium, they change shape and transition to an intermediate cellular state characterized by nuclear-driven cellular dynamics. The time-lapse microscopy images also indicate that no transient cellular protrusions were observed, as the round cells gradually acquired a neuron-like morphology (neuronal polarization). Furthermore, we also observed that the transformation of hBM-MSCs, hDP-MSCs and HDFa into neuron-like cells occurs rapidly and without cell division. It is important to note that previous studies have shown that the direct reprogramming of human somatic cells into neuronal cells also occurs rapidly and without cell division [9, 12, 15].

Therefore, our results provide further evidence that these intermediate cells, which differ in morphology and nuclear dynamics from both the original mesenchymal and fibroblast cells and the final neuron-like cells, may represent a key mechanistic stage in the transdifferentiation process. Further research will be needed to characterize these intermediate cells and determine whether they exhibit features of pluripotency or represent a distinct transitional phenotype, since, as we noted earlier, we have observed that intermediate cells can rapidly and repeatedly switch lineages without cell division [9, 12, 15].

It is important to note that, in our opinion, cells with lobulated or binucleated nuclei may also appear to be present in direct reprogramming experiments conducted using both the overexpression of transcription factors [9] (their Fig. S8B) as well as small-molecule-based protocols [18] (their Figs. 1D, S1D, S2A, S2B). Therefore, it would be interesting to examine whether these intermediate cells also appear in the various techniques described for cell reprogramming and transdifferentiation.

This work also highlights that neuron-like cells derived from human somatic cells exhibit neuronal polarization, develop a cytoskeleton remarkably similar to that of primary neurons, express specific neuronal and synaptic markers, form interconnected networks and exhibit plasticity. These findings confirm our previous results [26–29] and provide further evidence that hMSCs can be converted into functional, authentic neurons. As noted above, in this study we were unable to apply electrophysiological techniques that would have allowed us to conclusively demonstrate the successful generation of neurons derived from hBM-MSCs, hDP-MSCs, and HDFa. However, previous studies have shown that hDP-MSCs [43] and HDFa [10, 12, 14, 15, 18] can be differentiated into neuronal cells that exhibit functional excitability, as demonstrated by patch-clamp experiments. Futures studies incorporating functional assays will be essential to definitively confirm that hBM-MSCs display functional excitability.

As noted above, cell reprogramming and transdifferentiation techniques are characterized by their low efficiency, high complexity, and considerable random and stochastic behavior [16, 17, 19, 22, 23]. Small-molecule-based reprogramming protocols are generally considered a safer, more affordable, and more cost-effective alternative to traditional methods based on transcription factors [19]. Therefore, further efforts are needed to improve culture conditions and test different combinations of small-molecule cocktails in order to identify the key molecules required for effective conversion and to improve the survival and maturation of induced neurons. In this study and in previous ones, we showed that dibutyryl-cAMP and/or forskolin (an cAMP activator) can induce the conversion of hMSCs into neuron-like cells [28, 29]. It is important to note that dibutyryl-cAMP and/or forskolin are also used in many cell conversion protocols, such as indirect reprogramming, direct differentiation and direct reprogramming [12, 15, 16, 18, 19, 52]. Furthermore, our results also show that the most effective way to improve the survival of hMSC-derived neuron-like cells during transdifferentiation is to place a glass coverslip over the cells. Although further research is needed to understand the mechanisms underlying this phenomenon, previous studies have reported that hypoxia, a possible effect of the glass coverslip method, enhances cellular reprogramming [53] and may also promote the growth of neural stem cells and maintain their survival in vitro [54, 55]. Taken together, these findings may help us better understand not only which molecules are actually necessary for the effective conversion of human somatic cells into neurons, but also how to improve the survival of induced neurons, which would ultimately facilitate their use in regenerative medicine.

Finally, numerous authors have reported that the nuclei of many cultured hippocampal neurons [56] (their Fig. S2) and neural stem cells located in the ventricular-subventricular zone of the anterolateral ventricular wall of the human fetal brain [57] (their Fig. 2C) and adult mouse brain [58, 59] (their Figs. 1E, 4I, 6F, S1B, S2B, S6A and their Fig. 3A, respectively) as well as cells in the subcallosal zone [56], also have two lobes connected by an internuclear bridge. As noted in a previous study, we observed that the nuclear morphology of hBM-MSCs during neural transdifferentiation exhibits numerous similarities to the nuclear morphology described in the studies mentioned above [29].

Traditionally, adult neurogenesis has been thought to rely on active neural stem cells that continuously divide to create new neurons [60]. However, it is crucial to note that almost none of reports describing the presence of newly formed neurons in the adult brain have shown mitotic figures, which would confirm that adult neurogenesis occurs progressively through sequential phases of proliferation. Furthermore, the approaches used to investigate stem cell division and differentiation, and even direct lineage tracing, are inherently limited [61]. Moreover, recent studies have identified an alternative pathway in specific regions of the mammalian brain where neurogenesis can occur without cell division [62, 63]. Studies on direct neuronal reprogramming and our findings support the idea that neurogenesis can occur without cell division, calling into question the traditional view that neuron generation requires the asymmetric division of neural stem cells.

### Conclusions

In conclusion, our findings advance the understanding of the conversion of human somatic cells into neurons by providing new evidence that human somatic cells can adopt neural fate without cell division. Our results also further support the notion that the conversion of human somatic cells into neuron involves a transition through a transient intermediate state. Further research is needed to characterize these intermediate cells, which could prove crucial for elucidating the mechanisms underlying these cellular conversion processes and, ultimately, enabling their effective application in regenerative medicine.

## Declarations

### Ethics approval and consent to participate

The authors declare that all protocols used to obtain and process all human samples were approved by the local Ethics Committee of the University Hospital Virgen de la Arrixaca according to Spanish and European legislation and conformed to the Ethical Guidelines of the Helsinki Declaration. Donors provided written informed consent before obtaining samples.

For hBM-MSC experiments: (1) Title of the approved project: Validation of the collection of bone marrow mononuclear cells associated with the implementation of cell therapy clinical trials; (2) Name of the institutional approval committee or unit: Local Ethics Committee of the University Hospital Virgen de la Arrixaca; (3) Approval number: HCUVA10/13.25/11/2013; (4) Date of approval: 2013/11/25.

For hDP-MSC experiments: (1) Title of the approved project: Graphene-enhanced silk fibroin scaffolds for dental pulp regeneration (biographene); (2) Name of the institutional approval committee or unit: Local Ethics Committee of the University Hospital Virgen de la Arrixaca; (3) Approval number: HCUVA1.0/10/2022; (4) Date of approval: 2022/10/25.

### Consent for publication

Not applicable

### Availability of data and materials

All data generated or analysed during this study are included in this published article [and its supplementary information files].

### Competing interests

The authors declare that they have no competing interests.

### Funding

This work was supported by Instituto de Salud Carlos III (ISCIII) through the Spanish Network of Advanced Therapies Plus (TERAV+), RICORS subprogram, project RD24/0014/0001, co-funded by European Unión. The projects of the participating research groups are RD24/0014/0040 to M.B and RD24/0014/0022 to S.M.

### Author contributions

CB conceived of the study, designed the study, carried out lab work and drafted the manuscript; MM-M and FJR-L carried out lab work; DG-B carried out lab work and helped draft the manuscript; SM, JMM and MB helped draft the manuscript and provided financial support. All authors read and approved the final manuscript.

## Acknowledgements

We greatly appreciate the technical assistance of Microscopy Section Service of the University of Murcia.

## Notes

### Competing Interest Statement

The authors have declared no competing interest.

### Summary of Updates

In this revision of the article, the title, abstract, and content have been modified to facilitate the understanding and interpretation of the results obtained

